# Appearance of Amyloid-β Early in Life Initiates Neuronal Hyper-excitability, Mitochondrial Decay, and Loss of Dendritic Complexity in the Hippocampal CA1 Region of 5xFAD Mice

**DOI:** 10.64898/2026.02.04.703883

**Authors:** Amanda R. Kelley, Emily Sackinger, Mathew Frischman, Nicholas Thomas, Kaitlyn J. Kim, Grace Scuderi, Edwin M. Labut, Duncan MacMurchy, Toren Ikea-Mario, Jacob Rauenhorst, Easton Neitzel, Tacita Vu, Leda Liko, Alexandra Hoff, Ashley Hoff, Olivia Wallace, Wren K. E. Harry, Benjamin Hagen, Ken Lee, Alejandro Z. Bihun, Ibrahim A. Abou-Seada, Judy A. Butler, Fikru Nigussie, Arpa Ebrahimi, Phoebe Y. Lee, Luke C. Marney, Claudia S. Maier, Kathy R. Magnusson, Tory M. Hagen

## Abstract

Alzheimer’s disease (AD) is characterized by progressive cognitive decline and stereotyped neuropathology, yet the earliest cellular events that precede overt plaque burden and measurable behavioral impairment remain incompletely defined. Here, we tested the hypothesis that synaptic hyperexcitability and subcellular metabolic dysfunction emerge early in the 5xFAD mouse model and contribute to region-specific neuronal vulnerability before substantial amyloid plaque deposition.

Using the 5xFAD heterozygous mouse, we first established the onset of transgene expression and the timing of plaque accumulation. Robust transgene expression was detected by postnatal day 15 and significant plaque accumulation by 4 months of age. Ex vivo electrophysiology revealed an early hyperexcitable phenotype at 1 month of age, including both increased AMPA receptor-mediated transmission and N-methyl-D-aspartate receptor signaling associated with the GluN2B subunit. Given the tight coupling between glutamatergic hyperactivity, calcium dysregulation, and mitochondrial health, we assessed mitochondrial structure and function at this pre-plaque stage. Mitochondrial abnormalities consistent with impaired bioenergetic homeostasis were evident. Morphological analyses further demonstrated that these early changes were associated with altered dendritic architecture in the CA1 and dentate gyrus regions, revealing hippocampal subregional susceptibility. Finally, spatial transcriptomics supported this anatomical selectivity by identifying regionally enriched molecular signatures consistent with differential vulnerability. The CA1 region exhibited more reductions in mitochondria-related transcripts than CA3 or dentate gyrus and these reductions were specifically associated with CA1 pyramidal cell neurons.

Together, these findings define a pre-plaque window in 5xFAD mice marked by GluN2B-linked glutamatergic hyperexcitability, early mitochondrial disruption, and selective dendritic and transcriptional vulnerability across hippocampal subregions. This integrated timeline suggests that synaptic and metabolic dysfunctions arise before substantial plaque deposition and may represent tractable early targets for intervention aimed at delaying or preventing downstream neurodegeneration in AD.

## 1 Introduction

Alzheimer’s disease (AD) is a progressive neurodegenerative disorder that increasingly impairs memory and thinking skills, and robs the patient of the ability to carry out even simple tasks of daily living (National Institute on Aging, 2016). While the initializing events leading to the sporadic form of AD are still poorly understood (Mertas and Bosgelmez, 2025), AD ultimately affects brain regions important for memory and cognition, such as the hippocampus and neocortex (Fischer et al., 2025, Woodward et al., 2024, Rao et al., 2022). AD incidence is increasing in both the developed and developing worlds, and it is now estimated that 1 in 85 persons will be living with the disease by 2050, and 43% of those afflicted will not be able to meet normal activities of daily living without assistance. Currently, unpaid care-giving costs alone is estimated to be $346.6 billion for AD (Association, 2024). In the United States, at the time of this writing, an estimated 6.9 million are afflicted with AD and it is considered the 6th leading cause of death in America (National Institute on Aging, 2016, Association, 2017). There is no cure for AD, and current treatments are mainly palliative in nature (Buccellato et al., 2023, Sheikh et al., 2023).

There appears to be no single cause for the sporadic forms of AD; rather, a toxic stew of environmental, biological, and genetic factors are involved to varying degrees (Zhang et al., 2021, Mertas and Bosgelmez, 2025, Zhang et al., 2024, Suresh et al., 2023). Moreover, because one or more of these predispositional factors may occur decades prior to AD onset (Jack et al., 2013, Selkoe and Hardy, 2016, Aiello et al., 2025, Johansson et al., 2023, Caselli et al., 2020), definitive early hallmarks by which preventative strategies may be employed to mitigate risk for overt dementias have been equally daunting to discover. As such, chemopreventive strategies have been difficult if not impossible to implement. Despite these significant hurdles, a number of important biological markers for AD are now known.

At the neuro-anatomical level, neurite dystrophy, loss of dendritic complexity and spine loss in pyramidal neurons of the CA1 region of the hippocampus and cortex are consistently evident in post-mortem brains of AD patients (Perez-Cruz et al., 2011, Shi et al., 2020). Often this dystrophy is evident close to amyloid-β (Aβ) plaques (Blazquez-Llorca et al., 2017, Grutzendler et al., 2007), although they are also seen in tauopathies that lack amyloid plaques (Shi et al., 2020). It is unknown if these changes occur early in the disease process or only after amyloid plaque formation

In addition to anatomical alterations, the signature pathologies of AD are amyloid plaques and neurofibrillary tangles (Mirra et al., 1991, Armstrong, 2006). These lesions are evident in both sporadic and familial forms of AD (Liu et al., 2024, Braak and Braak, 1991). Familial and early onset AD has a strong genetic component and many cases are associated with mutations in presenilin (**PSEN**) and amyloid precursor protein (**APP**) genes (Wu et al., 2012). PSEN is a subunit of gamma-secretase, which processes APP to Aβs of 38-43 amino acids (Esler and Wolfe, 2001, Takami et al., 2009). Mutations of PSEN and APP tend to produce more Aβ42 than in non-transgenic mice (Citron et al., 1998, Eckman et al., 1997). There is evidence that enhanced levels of amyloid, tau, and brain shrinkage begin to increase 2 to 3 decades before evidence of dementia in individuals who will eventually develop AD (Masters et al., 2015, Bateman et al., 2012, Ringman et al., 2012, Ringman et al., 2008). This suggests that prevention strategies for AD should, if possible, focus on early events in disease pathology or, most importantly, in disease initiation.

Based on the aforementioned hallmarks of AD, a number of transgenic animal models have been developed which replicate overt amyloidosis and/or tau pathologies associated with AD. The most common transgenic models exploit overexpression of amyloid, which is the basis of the amyloid hypothesis of AD (Selkoe and Hardy, 2016). These models show, to varying degrees, common attributes of pathologies associated with the sporadic human form of the disease (Preuss et al., 2020). They can be useful for studying the timeline of disease progression.

Multiple strains of PSEN mutant mice show a transient increase in N-methyl-D-aspartate receptor (NMDAR) transmission (as shown by input/output curves) at an early age (3 months), followed by deficits later in life (Auffret et al., 2010, Auffret et al., 2009, Dewachter et al., 2008, Wang et al., 2009). This has been shown to be associated with an increase in NMDA receptor GluN2B subunit expression in the PS1A246E PSEN mutant (Dewachter et al., 2009). Glutamate receptors, such as the NMDARs and α-Amino-3-hydroxy-5-methylisoxazole-4-propionic acid hydrate (AMPAR) receptors are in high density in AD-affected brain regions, such as hippocampus and cortex (Magnusson, 1998, Magnusson and Cotman, 1993). Although NMDARs normally initiate memories and synaptic plasticity (Morris et al., 1986, Alessandri et al., 1989, Butelman, 1989, Heale and Harley, 1990, Mondadori et al., 1989), overstimulation by glutamate can lead to calcium overload-related excitotoxicity and death of cells (Choi, 1992). As the PSEN mutants do not develop the hallmark pathologies (Vidal et al., 2012, Janus et al., 2000), it is not clear whether the NMDAR hyperactivity contributes to the development of these pathologies.

A commonly used transgenic mouse strain that exemplifies many of the common pathologies of AD, 5XFAD transgenic mice (**5XFAD**), were created by Oakley and coworkers (Oakley et al., 2006). They express 2 PSEN and 3 APP mutations (Citron et al., 1998, Eckman et al., 1997, Goate et al., 1991, Mullan et al., 1992) and overexpress Aβ42 (Eimer and Vassar, 2013). This model shows more significant positive correlations with late-onset human AD in gene expression for the immune and neuronal systems, cell cycle and stress than many other mouse models (Preuss et al., 2020). They do develop amyloid plaques, unlike PSEN mutants (Vidal et al., 2012, Janus et al., 2000), but it is currently believed that this occurs before any synaptic dysfunctions. Interneuronal Aβ42 is identifiable at 1.5 months of age and plaque formation and gliosis occur by 2 months (Eimer and Vassar, 2013, Oakley et al., 2006). Declines in total NMDAR fEPSP synaptic transmission input/output and early LTP are seen at 6 months of age, but these measures are equivalent in transgenic and WT mice at 3.5-4.5 months (Kimura and Ohno, 2009). Neuronal loss also becomes detectable by 6 months (Eimer and Vassar, 2013). Cognitive deficits have been seen by 3 to 6 months of age in 5xFAD Het mice (Jawhar et al., 2012, Oakley et al., 2006), although one research group reports spatial memory deficits in the Morris water maze as early as 1 month of age (Tang et al., 2016, Wu et al., 2018). Mitochondrial dysfunction, based on RNA-SEQ changes, however, shows up as early as 7 weeks (Kim et al., 2012). Given that 5XFAD mice express 2 PSEN mutations, it seems likely that they might show early transient NMDAR hyperexcitability and consequences of increased cytosolic calcium values.

NMDARs conduct Ca^2+^ and there is an abnormal rise in internal Ca^2+^ in PSEN mutant mice (Schneider et al., 2001), which could potentially involve Aβ and GluN2B subunits (Ferreira et al., 2015). This rise was attributed to alterations in storage of calcium in the endoplasmic reticulum (Schneider et al., 2001), however, it could also be due to mitochondrial dysfunction (Stanika et al., 2009, Ma et al., 2020, Ferreira et al., 2015, Ryan et al., 2020, Mustaly-Kalimi et al., 2025, Walters and Usachev, 2023).

Impaired energy metabolism is one of the earliest and most consistent features of AD (Yu et al., 2022, Mosconi, 2013). This reduction in energy metabolism is closely linked to mitochondrial dysfunction. Molecular studies further highlight mitochondrial abnormalities in AD, including variations in size, structural changes, and impaired expression of nuclear-encoded respiratory chain components. Electron microscopy has revealed specific mitochondrial alterations in AD brains, such as size heterogeneity, disrupted cristae, accumulation of osmophilic material, presence of lipofuscin vacuoles, and elongated, interconnected organelles. Studies showing significantly decreased antioxidant levels and altered expression of antioxidant enzymes in AD brains, as well as oxidative stress markers that are closely linked to significant loss of synaptic proteins in the brains of MCI and pre-AD patients indicate that mitochondrial dysfunction plays a central role in AD pathology.

The current study addressed the hypothesis that there are early subcellular changes in the 5xFAD mouse model that occur before significant amyloid plaque deposition or reported cognitive deficits. This study determined the age of onset of significant transgene expression by 15 days of age and plaque accumulation at 4 months old in the 5xFAD mouse model. Electrophysiological recording confirmed that 5xFAD Hets exhibited hyperexcitability of both AMPAR and NMDAR subunit GluN2B at 1 month of age. Since calcium dysregulation can be influenced by dysfunction of mitochondria (Ryan et al., 2020, Mustaly-Kalimi et al., 2025, Walters and Usachev, 2023), mitochondrial morphology and function at this young age were examined. The impact of these changes on dendritic morphology revealed a hippocampal subregional susceptibility, which was substantiated by spatial transcriptomics.

## 2 Materials and Methods

### 2.1 Animal husbandry

The 5xFAD mouse model used for this research project, B6.Cg Tg(APPSwFlLon,PSEN1*M146L*L286V)6799Vas/Mmjax, RRID:MMRRC_034848-JAX, was obtained from the Mutant Mouse Resource and Research Center (MMRRC) at The Jackson Laboratory, an NIH-funded strain repository, and was donated to the MMRRC by Robert Vassar, Ph.D., Northwestern University. 5xFAD Het (MMRRC/JAX) male mice were bred with female C57BL/6J mice (JAX Labs, Bar Harbor, ME) to produce non-transgenic littermates (referred to as wildtype (**WT)**) or 5xFAD heterozygous (**5xFAD Het**) mice. Mice were fed *ad libitum* with standard rodent chow and kept on a 12-hour light/dark cycle and housed in AALAC-approved space under veterinary supervision at Oregon State University. Experiments were performed according to protocols approved by the Oregon State University Institutional Animal Care and Use Committee (IACUC-2019-0043, IACUC-2020-0082, ACUP4921). Mice used for EM, Golgi analysis and spatial transcriptomics were sacrificed using isoflurane and decapitation, and brains were quickly removed for processing. Mice used for electrophysiology and immunohistochemistry were euthanized as described below for electrophysiology. Non-transgenic littermates (WT) and 5xFAD Het mice were collected at 15 days or 1, 2, or 4 months of age for immunohistochemistry and at 1 month of age for all other studies. Ns for each experiment are indicated in the figure legends.

### 2.2 Genotyping

When mice reached approximately 15 days of age, ear punches were collected for identification and genotyping purposes. Genotyping was performed either by Transnetyx, Inc. (Cordova, TN) or in-house. Locally, ear punch samples were digested overnight in a GNT buffer containing proteinase K at 37°C. After digestion, insoluble materials were centrifuged and pelleted. The supernatant was collected and used for PCR amplification using Promega GoTaq Master Mix (Promega, Cat. # M7122). Primer sequences were provided by Jackson Laboratories and are shown below. A thermocycler with the following five-step protocol: 1) 94°C for 2 minutes, 2) 95°C for 30 seconds, 3) 65°C for 1 minutes 30 seconds with temperature decreasing by 0.5°C each cycle, 4) 68°C for 1 minute, 5) repeat steps 2-4 for 10 cycles total (touchdown), 6) 94°C for 30 seconds, 7) 60°C 1 minutes 30 seconds, 8) 72°C for 1 minute, 9) Repeat steps 6-8 for 28 cycles total, 10) 72°C for 5 minutes, and 6) 4°C hold. Following the completion of the PCR amplification, samples were run on an SDS-polyacrylamide gel using Laemmli buffer, and bands were visualized using a handheld UV light. WT animals have a band at 219bp only, and 5xFAD Het animals have an additional band at 129bp.

#### Genotyping Primers

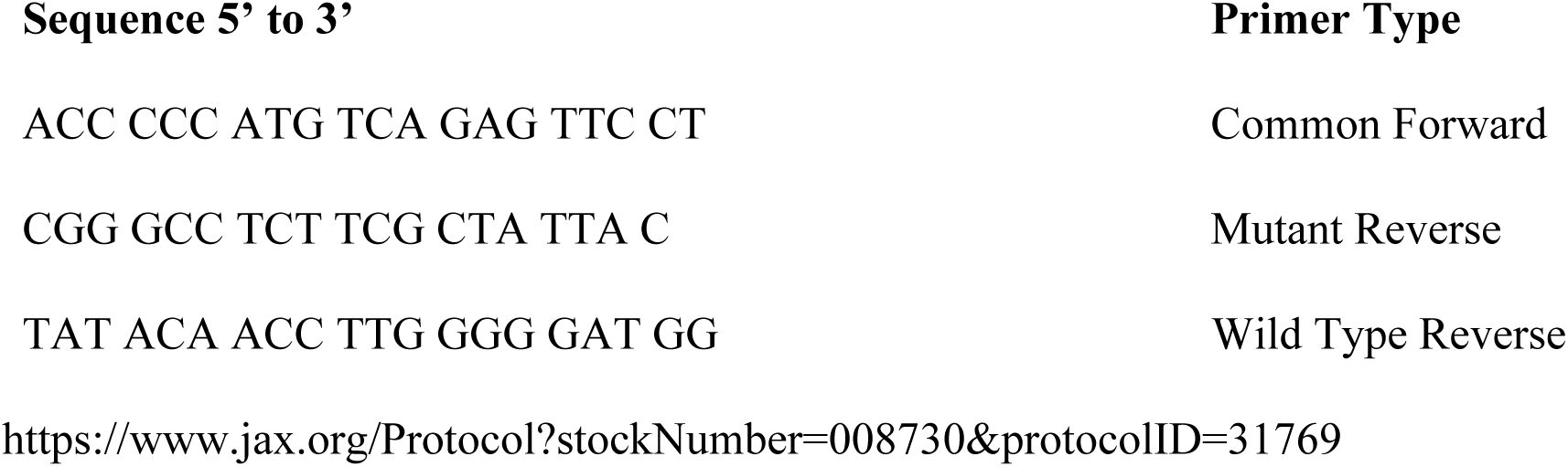

### 2.3 QPCR: Transgene expression

To assess the onset of mRNA expression of the transgene, small pieces of brain tissue, weighing approximately 10 mg, were used to perform QPCR on WT and 5xFAD Het animals. A total of 6 animals (male & female combined) were used for each group analyzed. Total RNA was extracted using the Qiagen RNeasy Lipid Tissue Mini Kit (Cat. # 74804), and RNA concentration was measured using a Nanodrop. Following extraction, 20ug of total RNA was used to create cDNA using a Thermo Fisher SuperScript IV kit (Cat. # 18091050). Samples were prepared for QPCR analysis using TaqMan Gene Expression Master Mix (Cat. # 4370048), and expression was measured using a StepOne Plus RT-PCR machine. All samples were run using the following setup: 1) 50°C for 2 minutes, 2) 95°C for 10 minutes, 3) 95°C for 15 seconds, 4) 60°C for 1 minute, 5) Repeat steps 3-4 for a total of 40 cycles, 6) 4°C hold. TaqMan primer/probe mix was obtained for GAPDH (Assay ID, Mm99999915_g1). Gene expression for TaqMan primers/probes was obtained using standard ΔΔct calculations. Transgene expression was obtained using custom ordered primer/probes from Sigma with sequences provided by Jackson Laboratories (shown below). Due to significant differences in primer efficiency between the transgene primer/probe set and TaqMan GAPDH primer/probe set, the Pfaffl method was used to calculate relative abundance. For transgene expression, all samples were normalized to cortical tissue from a 0.5 month old WT animal.

#### QPCR 5xFAD Transgene Primers and Probe

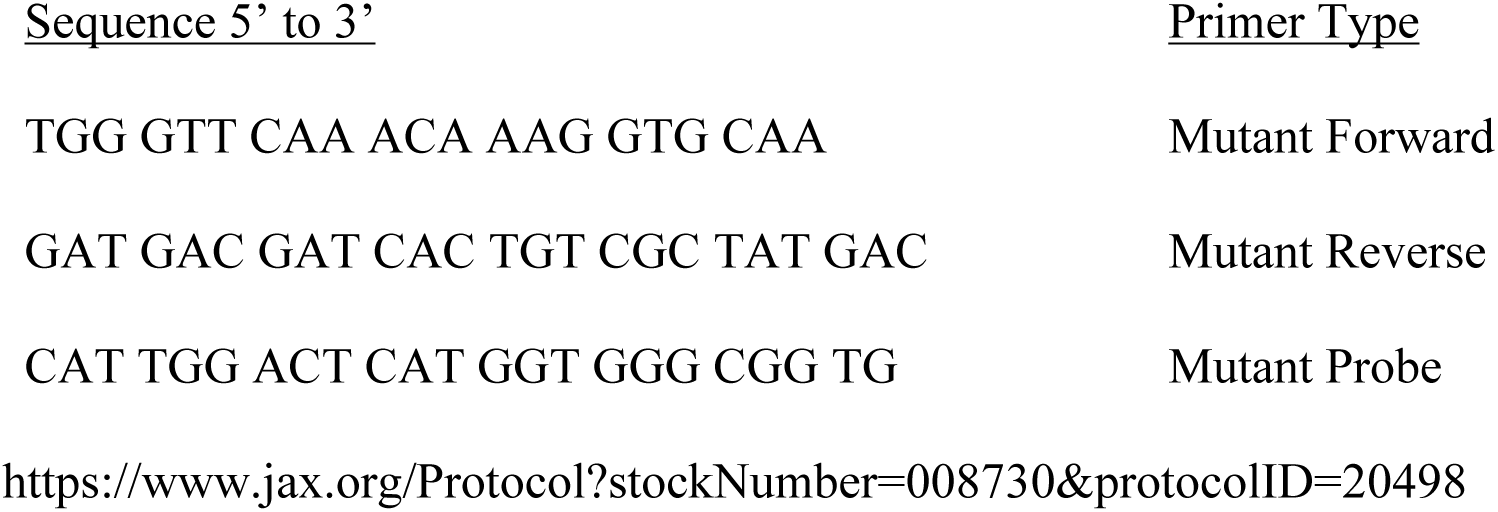

### 2.4 Immunohistochemistry

Immunohistochemical staining for beta-amyloid was carried out on hippocampal slices from 0.5-, 1-, 2-, and 4-month-old male and female WT and 5xFAD mice (see euthanasia below) with the following steps: Hippocampal slices were: 1. washed 3 X 10 minutes with stirring with Phosphate Buffered Saline with Tween 20 (PBS-T: pH ≈ 7.2, 0.04 M Na_2_HPO_4_, 0.01 M NaH_2_PO_4_ᆞH_2_O, 0.14 M NaCl, 0.1% Tween 20), 2. immersed for 20 min in 0.3% hydrogen peroxide in PBS to inhibit endogenous peroxidase in red blood cells prior to staining, 3. blocked for 1 hour with 3-5% goat serum in PBS with 0.1% Triton X and 0.02% sodium azide, 4. incubated overnight with 1:500 rabbit anti-beta amyloid antibody (abcam ab2539 rabbit polyclonal aby) at room temperature, and 5. rinsed in PBS + Tween 20 3 X 10 minutes, 6. incubated in biotin-conjugated secondary antibody (1:500 goat anti-rabbit IgG antibody) with 5% goat serum in PBS with 0.1% Triton X and 0.02% sodium azide for two hours, 7. rinsed in PBS + Tween 20 3 X 10 minutes, 8. incubated with 1:800 Vectastain Avidin-Biotin Complex Kit (ABC: Vector Laboratories; Burlingame, CA) in PBS for 1 hour at room temperature, 9. washed with Tris buffered saline with tween (TBS-T: pH ≈ 7.5, 0.005 M Tris Base, 0.0038 M Tris HCl, 0.015 M NaCl, 0.1% Tween 20) 3 X 10 minutes, and 10. incubated in 3,3′-diaminobenzidine (DAB) solution plus hydrogen peroxide (SIGMAFast, D4293, Sigma-Aldrich, St. Louis, MO) for 7 minutes to form a brown precipitate over the protein of interest, beta amyloid. Following 3 X 10-minute rinses in TBS-T, hippocampal slices were mounted onto slides and dehydrated using 70%, 95%, and 100% ethanol and cleared with xylene for 5 minutes each prior to adding a coverslip with Mount-Quick mounting media (Daido Sangyo CO. LTD, prod #19478, Japan). For each individual mouse, 200X images were acquired from three different areas within the CA1 stratum radiatum of the hippocampus from 3 different slices, for a total of nine images per mouse, with the use of a Nikon inverted microscope. All neuronal images were analyzed using NIH ImageJ software, with a maximum brightness level of 170 and a threshold of 70 using the default thresholding macro. The percent area of the image with pixel intensity above the threshold was determined. If there were any gross artifacts present on the image, the lasso tool was used to trace around the artifact. All image analyses were performed with the analyst blind to the treatment, genotype and sex of the mouse. Statistical analysis was performed via three-way analysis of variance (ANOVA, p < 0.05) with the use of Statview software.

### 2.5 Electrophysiology

For electrophysiology and immunohistochemistry experiments, 1 month old female WT and 5xFAD Het mice were anesthetized via intraperitoneal injection with 100 mg/kg ketamine and 20 mg/kg xylazine, followed by exposure to isoflurane. Animals were then rapidly perfused transcardially, with ice-cold carboxygenated artificial cerebrospinal fluid (aCSF; NaCl (124mM), KCl (2mM), KH2PO4 (1.25mM), MgSO4 (2mM), Glucose (10mM), NaHCO3 (26mM)), with no added Ca^2+^). Following removal of the brain, a vibratome was used to obtain 300µm coronal brain slices from freshly isolated brain tissue in WT or 5xFAD mice, keeping all tissue submerged in ice cold carboxygenated aCSF. After sectioning, slices were allowed to recover in a slice keeper containing aCSF with 2mM Ca^2+^ at 32°C for 30 minutes followed by a 2.5 hour incubation at room temperature. After recovery, slices from 1 month old WT or 5xFAD Het mice were placed onto a 16-electrode Med 64 Quad II probe (AutoMate Scientific, Cat. # MED-PG501A). The stratum radiatum of the CA1 region of the hippocampus was positioned over the electrodes. Slices were perfused with carboxygenated aCSF at 2mL/min at 32℃. Schaffer collateral axons from the CA3 neurons were stimulated, and field excitatory postsynaptic potentials (fEPSP) from neurons in the CA1 region were recorded with the use of the MED 64 system and Mobius software (AutoMate Scientific, Berkeley, CA).

Baseline recordings were performed until fEPSP was stable for at least 10 minutes. The contributions of specific receptors or receptor subunits were determined with the use of sequential addition of antagonists against AMPARs (6,7-Dinitroquinoxaline-2,3-dione (DNQX), 30µM), gamma-aminobutryic acid receptors (GABAR, Pictrotoxin, 10µM), and GluN2B (Ro 25-6981, 4µM; Fig. 3A). After each drug administration, the fEPSP was allowed to stabilize for at least 20 minutes and an input (fiber volley)/output (fEPSP; I/O) curve was obtained. An I/O consisted of stimulations from 10-60 pAmps in 5 pAmp increments every 20 seconds. Each stimulation level from post-treatment I/O curves were subtracted from the same stimulation level from a prior I/O curve, and the differences were then expressed as the I/O curve for a specific receptor or subunit component of the fEPSP (Fig. 3B). Statistical analysis using an extra sum-of-squares F Test comparison of non-linear regression curve fits for the drug study differences I/O were performed with Prism software.

### 2.6 Electron Microscopy

#### 2.6.1 Animals and Tissue Collection

Heterozygous 5xFAD mice (N = 6) and wild-type (WT) littermates (N = 3) at one month of age were used for all experiments. All procedures were performed in accordance with institutional and NIH guidelines for the care and use of laboratory animals. Mice were the control group of a treatment study and received PBS injections IP every 3 days from post-natal day 15 to 30. Mice were euthanized as described previously, and whole brains were rapidly removed. Hippocampi were dissected and immediately immersed in modified Karnovsky fixative at 25 °C until further processing by the Oregon State University Microscopy Center.

#### 2.6.2 Tissue Processing and Post-Fixation Staining

Following fixation, hippocampal samples were embedded in agarose to facilitate post-fixation staining. Embedded tissues were rinsed in 0.1 M sodium cacodylate buffer and subsequently treated with 1.5% (w/v) potassium ferrocyanide and 2% (w/v) osmium tetroxide in distilled water (T-O-T-O staining). Samples were then incubated in uranyl acetate and lead aspartate to enhance contrast.

#### 2.6.3 Dehydration and Resin Embedding

Tissues were dehydrated through a graded ether–acetone series (10%, 30%, 50%, 70%, 90%, 95%, 100%, and 100%), with each step lasting 10–15 min. After complete dehydration, samples were infiltrated with and embedded in araldite resin.

#### 2.6.4 Sectioning and Electron Microscopy

Approximately 30 resin-embedded sections were prepared from each hippocampus and further sectioned into ultrathin slices (∼5 µm thick). Ultrathin sections were imaged using a FEI Helios Nanolab 650 electron microscope operated in scanning transmission electron microscopy (STEM) mode. Images containing synaptic structures were selected based on the presence of synaptic vesicles, post-synaptic electron densities, and characteristic “railroad track” double membranes indicating synaptic boundaries.

#### 2.6.5 Image Selection and Quantitative Analysis

Images that met these structural criteria were analyzed for mitochondrial number and morphology. Mitochondrial counts were performed manually on each selected image. Mitochondrial size heterogeneity was quantified using **FIJI** (ImageJ) software. Cristae integrity was evaluated independently by two blinded investigators using a semi-quantitative scoring system. Mitochondria with fully intact cristae were assigned a score of **1**; those showing partial cristae disruption or denudation but retaining some intact structures were scored as **2**; and mitochondria displaying widespread cristae collapse or near-complete loss of structure were scored as **3**. Representative examples of each category are provided in Figure 4.

### 2.7 Golgi Stain and Analyses

#### 2.7.1 Golgi-Cox Rapid Stain

Whole brains were quickly removed from 1 month old WT and 5xFAD Het (male and female combined) mice, rinsed with deionized water, and stained using the Golgi-Cox Rapid Fast Stain Kit (FDNeurotechnologies, Cat. # PK401). Following the staining protocol, brains were gradually frozen in 2-methylbutane immersed in liquid nitrogen before being hard frozen on dry ice. Brains were stored at -80°c and sectioned at 100μm/slice by FDNeurotechnologies.

#### 2.7.2 Sholl Analysis, Neuron Length, and Area Under the Curve

To assess dendritic branching, 100μm Golgi-Cox stained tissue sections were used for the analysis of the CA1, DG and CA3 regions of the hippocampus. Totals of 30, 24, and 30 neurons were used for each group from CA1, CA3, and DG regions, respectively. Z-stack images were taken on a Leica DM6000 with the same light settings for each image. The objectives 20x, 10x, and 40x were used to image neurons from the CA1, CA3, and DG regions, respectively. Following capture of images, z-stacks were reassembled, converted to 8-bit image files, inverted, and adjusted for background noise with FIJI software (Schindelin et al., 2012). The Simple Neurite Tracer plugin (Longair et al., 2011) was used to trace the reassembled z-stacks. The Sholl analysis tool within the Simple Neurite Tracer plugin calculated the number of intersections, or dendritic branches, every 20μm from the soma. Additionally, the Simple Neurite Tracer plugin measured the length of every dendritic branch which was used to calculate total neuron length. Sholl analysis results were also used to calculate area under the curve using a formula for the area of a trapezoid. Analysis of images was performed blind.

#### 2.7.3 Spine density Analysis

To measure spine density, a static planar image from each hippocampal region (i.e. CA1, CA3, and DG) was taken on a Leica DM6000 with the 60x objective lens, and an in-focus dendritic segment was used for analysis. A total of 30 neurons were used to measure spine density in each group for CA1, CA3, and DG hippocampal regions. Apical spine density was measured for CA1 and CA3 neurons, but basal spine density was measured for DG neurons. The images were converted into 8-bit image files, inverted, cropped and adjusted for background noise with FIJI (Schindelin et al., 2012). The dendritic branch was turned into a binary and skeletonized image, and each protruding spine was counted manually. The dendrite length was measured using the Simple Neurite Tracer plugin (Longair et al., 2011). The number of spines and dendrite length were recorded, and spine density was calculated using an Excel spreadsheet. All spine density measurements were performed blind.

### 2.8 Spatial Transcriptomics

#### 2.8.1 Animals

Four one-month-old 5xFAD Het (C57BL/6 background) and four WT littermates (male and female combined) were raised from breeders. Mice were euthanized via CO_2_-induced narcosis followed by decapitation. Brains were swiftly removed and fresh-frozen with dry ice, then stored in a -80 °C freezer until further processing.

#### 2.8.2 Sample Processing

Cryostat sections (6 µm) through the dorsal hippocampus were mounted onto slides and shipped to Nanostring, Inc. (Seattle, WA), where they were treated with indexing, UV-photocleavable oligo-labeled cDNA probes using the *GeoMx Mouse Whole Transcriptome Atlas Mouse RNA for Illumina Systems* (Catalogue# GeoMx NGS RNA WTA Mm). The mRNA transcript counts were measured separately within the cell body layers of the CA1, CA3, and DG hippocampal subregions of each mouse. Brain sections were stained for NeuN, GFAP, Iba1, and DNA, and subregions were marked on scans. UV light was used to collect oligo-barcodes within each subregion, which were quantified with nCounter (Nanostring, Inc).

#### 2.8.3 Quality Control

Samples, separated into subregions, were grouped via t-SNE (van der Maaten and Hinton, 2008) or UMAP (McInnes et al., 2020) for data visualization. One Het mouse had no transcript detection, so it was excluded from analysis, giving WT n=4 and Het n=3.

#### 2.8.4 Differential Gene Expression Analysis

Raw counts were analyzed for differential gene expression by genotype in each individual subregion using NOISeqBio (Tarazona et al., 2015). Global counts-per-million (CPM) profiles from each sample were visualized and used to filter out transcripts with a CPM less than 10. Raw counts were normalized using a Trimmed Mean of the M-values (TMM) approach. Analysis used 1000 permutations for comparisons. NOISeqBio uses a ‘probability’ as a statistical measure that is equivalent to 1 – the False Discovery Rate (FDR) adjusted p-value. For simplicity, the probability was converted to an adjusted p-value by subtracting the probability from 1 (p = 1-probability). Genes were determined to be differentially expressed given an absolute value of the fold change greater than 1.5 (|FC|>1.5) and p < 0.05.

#### 2.8.5 Gene Set Enrichment Analysis

Gene Set Enrichment Analysis (GSEA) was performed using the GSEA software from UC San Diego and the Broad Institute (Mootha et al., 2003, Subramanian et al., 2005) and the Molecular Signatures Database (Liberzon et al., 2015, Liberzon et al., 2011). Genotype differences in Reactome (Gillespie et al., 2022) and Gene Ontology (Ashburner et al., 2000, Gene Ontology et al., 2023) database pathways were analyzed in each subregion with gene set permutation, using 1000 permutations for each comparison. Gene sets were determined to be differentially expressed given FDR <0.25.

*Data Visualization:* Data were visualized with Matplotlib (Hunter, 2007) and ggplot2 (Wickham, 2016).

#### 2.8.6 Spatial Transcriptomic Deconvolution and Pathway Analysis

To resolve cell-type specific alterations within the hippocampal formation, raw spatial transcriptomic counts were deconvoluted using the NanoString SpatialDecon algorithm (Danaher et al., 2022). Cell type abundance was estimated using a single-cell RNA sequencing reference derived from the adult mouse brain (Yao et al., 2021), enabling the enumeration of discrete neuronal (CA1, CA3, DG) and glial populations.

Differential expression between Heterozygous (HET) and Wild-Type (WT) genotypes was calculated for each cell type using a reverse deconvolution linear regression model. To assess metabolic function, we curated a targeted panel of mitochondrial genes involved in transport, biogenesis, and oxidative phosphorylation. Global shifts in this gene set were analyzed using a Chi-Square test of independence to determine if specific cell populations exhibited non-random regulatory patterns. Finally, functional biological impact was assessed via Gene Set Enrichment Analysis (GSEA) (Korotkevich et al., 2019), ranking genes by their cell-type-specific log2 fold change to identify coordinated regulation of Gene Ontology (GO) biological processes, with significance defined as an adjusted p-value < 0.25.

### 2.9 Experimental Design and Statistical Analyses

Statistical analyses were performed using GraphPad Prism 9, R studio, and G*Power. QPCR gene expression was analyzed by 2-way ANOVA and Fisher’s LSD post-hoc analysis. Electrophysiology data were analyzed using a non-linear regression curve fit and extra sums-of-squares F test to assess differences between genotypes and sexes. EM analysis used unpaired Welch’s t-test. For dendritic analyses, neurons were pooled and used to perform all statistical analyses. Sholl analysis data was analyzed using repeated measures ANOVA with post hoc comparisons and multiple testing corrections. β-amyloid plaque deposition and transgene expression, neuronal length, area under the curve, and spine density were analyzed using ANOVA and post-hoc analysis where indicated. All graphs and figures were made using GraphPad Prism.

## 3 RESULTS

3.1 **5xFAD transgene was detectable at 0.5 months of age**

There was a significant main effect of Age (p < .0001) and Genotype (p<.0001) and a significant Age X Genotype interaction (p < .0001) on transgene expression (Fig. 1). The transgene expression was significantly higher at 0.5 (p=.01) and 1 (p<.0001) month of age in the 5xFAD Het mice than in WT littermates (Fig. 1). The 5xFAD Het mice at 1 month of age also had significantly higher transgene expression than 0.5 month old 5xFAD Hets (p<.0001; Fig. 1).

**Figure 1.**
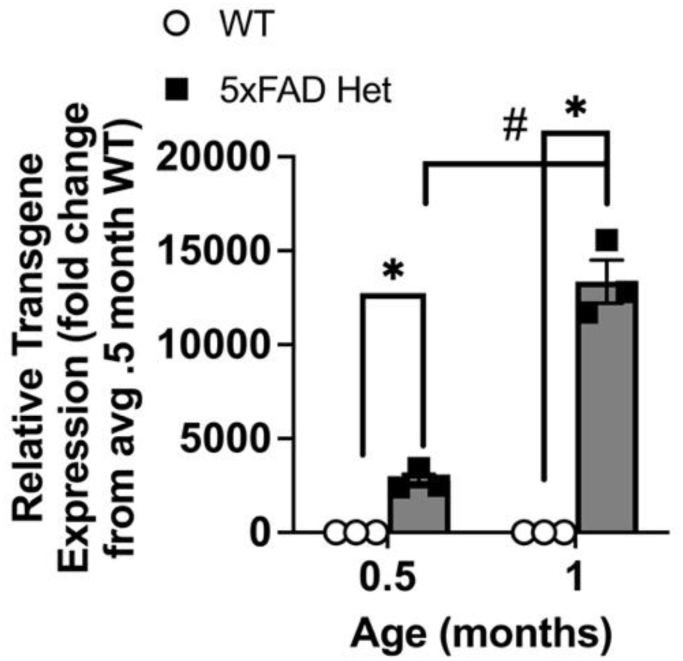
Measurement of 5xFAD transgene expression by qPCR in enriched CA1 tissue from 0.5 and 1 month old WT and 5xFAD Hets. There was significantly more transgene expression in 0.5 and 1 month old 5xFAD Het mice than in WT of the same age. ANOVA and Fisher’s post-hoc test. N = 3-4. Symbols indicate p<.05 for difference from WT (*) or for Hets between ages (#).

### 3.2 β-amyloid staining was not significantly increased until 4 months of age

There was a significant main effect of Age (p=.0059) and Genotype (p=.0067), but not Sex (p=.55), on percent area of β-amyloid immunostaining (Fig. 2). There was also a significant Age X Genotype X Sex interaction (p=.02). 5xFAD Hets had significantly more percent area of staining than WT over all ages and within the 4 month olds (p=.0001; Fig. 2D) with data collapsed across sex. Female 4 month old Hets also had more area stained than female 4 month old WT (p<.0001; Fig. 2D). The 2 and 4 month old mice had larger percent area of immunostaining than 1 month olds (p=.038-.049) and near significantly greater than .5 month olds (p=.059-.078; Fig. 2D).

**Figure 2.**
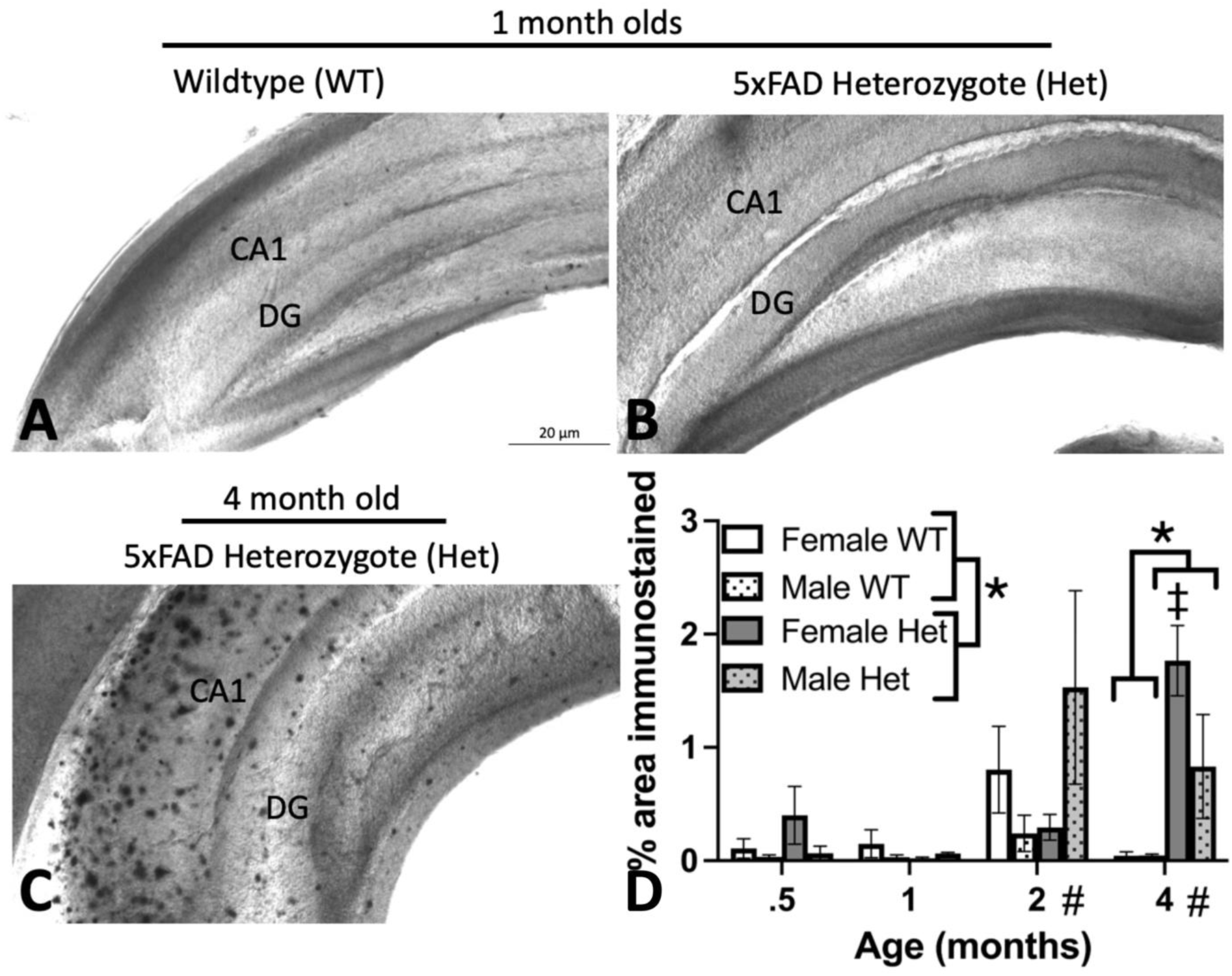
Plaque formation in the CA1 region of the hippocampus. Immunohistochemistry was performed to measure plaque formation with a β-amyloid antibody at .5, 1, 2, and 4 months of age. Representative images are shown for 1 month old WT (A) and 5xFAD Het (B) and 4 month old 5xFAD Het (C) mice. D) 5xFAD Het mice showed greater β-amyloid immunostained % area overall ages (*) and within the 4 month olds (*) than WT. Female Hets had a larger % area of staining than WT at 4 months of age (‡). Mice at 2 and 4 months had more % area stained than 1 month olds (#). ANOVA and Fisher’s LSD. N = 6-10. Symbols indicate p<.05 for difference.

### 3.3 Glutamatergic hyperactivity was present at 1 month of age

Glutamate receptor function was assessed to determine whether 5xFAD mice, which express two PSEN1 mutations, show NMDA receptor hyperactivity. With the use of hippocampal slices from 1 month old female WT and 5xFAD Het mice and a multielectrode array probe, slices were stimulated over the Schaffer collateral axons and recorded in the stratum radiatum of the CA1 region. Slices were exposed to different selective inhibitory drugs in order to determine the relative contribution of AMPA receptors or subunits of NMDA receptors (Fig. 3A-C). Female 5xFAD Het animals exhibited higher AMPA receptor (Fig. 3D; p<.0001) and GluN2B subunit (Fig. 3E; p = .024) responses for a given fiber volley input than WT at 1 month of age.

**Figure 3.**
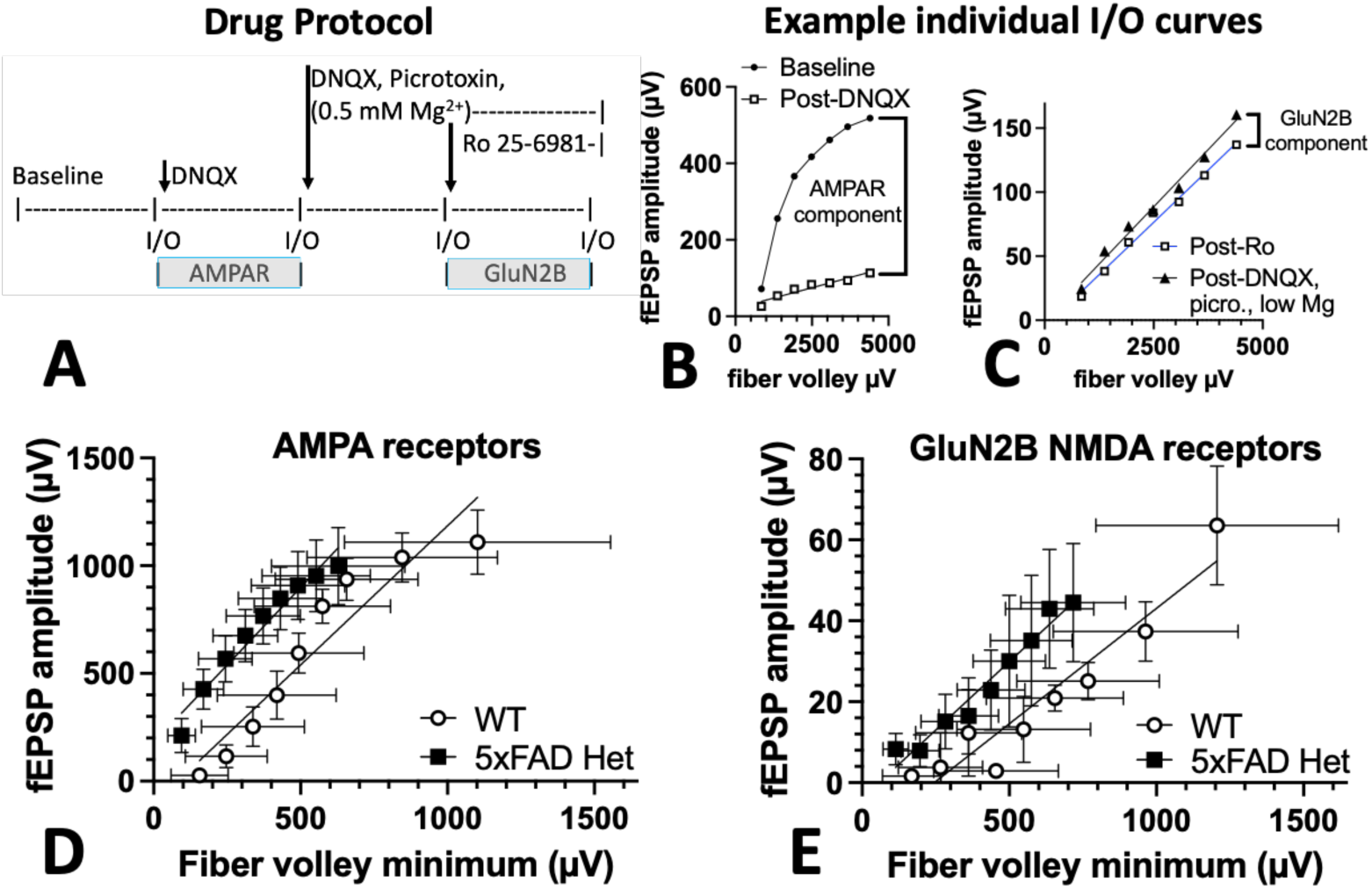
Genotypic differences in glutamate receptor activity at 1 month of age in CA1 region of hippocampal slices. A) Timeline of Input/Output sessions (I/O) and drug delivery. Differences between specific I/O curves for specific receptor components is indicated below the timeline. B,C) Example individual experiment plots of I/O curves post-drug stabilization for the AMPA receptor (B) and GluN2B subunit (C) components of the fEPSP. Brackets on the right indicate which curves were subtracted to obtain the receptor components. C,D) Female 5xFAD Het animals exhibited higher AMPA receptor (C) and GluN2B subunit (D) responses for a given averaged fiber volley input than wild type at 1 month of age. Extra sum-of-squares F Test comparison of non-linear regression curve fits. N=4-7. Symbols=mean, error bars = SEM.

### 3.4 Mitochondrial damage was increased at 1 month of age

Ultrastructural analysis of hippocampal synaptic mitochondria was performed using scanning transmission electron microscopy (STEM) in one-month-old 5xFAD heterozygous mice and age-matched wild-type (WT) littermates. For both genotypes, only mitochondria localized to synaptic regions—identified by the presence of synaptic vesicles, post-synaptic densities (PSD), and characteristic “railroad track” membranes—were included in the analysis (Fig. 4A,B). In WT mice, synaptic mitochondria exhibited uniform spherical morphology, consistent size, and homogeneous electron density. Cristae were densely packed and extended throughout the inner mitochondrial matrix, indicating intact ultrastructural organization (Fig. 4A). In contrast, hippocampal mitochondria from 5xFAD mice displayed pronounced abnormalities, including irregular shapes, variable size, and increased membrane electron density (Fig. 4B). These features were frequently accompanied by fragmented or vacuolated mitochondria, consistent with early degenerative changes. Despite these alterations, the overall number of mitochondria per synaptic image field did not differ significantly between genotypes (*t*(6.177) = .000, *p* >.9999, Welch-corrected unpaired two-tailed *t*-test; not shown).

**Figure 4.**
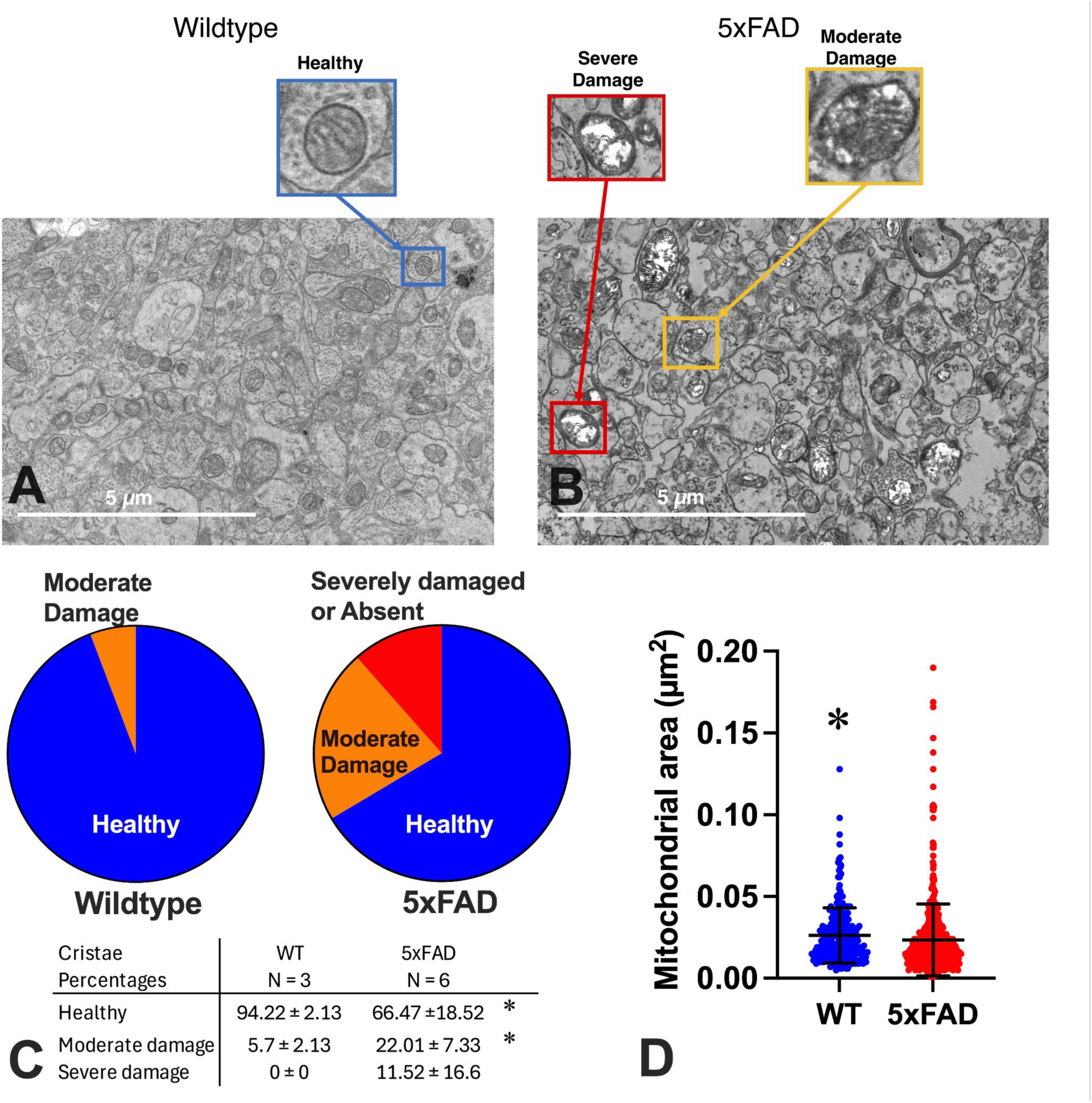
Representative electron micrographs and quantitative assessment of mitochondrial cristae integrity in hippocampal synapses of wild-type (WT) and 5xFAD heterozygous mice. A) Representative image of a hippocampal synaptic region from a WT mouse showing mitochondria with intact cristae and preserved synaptic structures, including synaptic vesicles and post-synaptic densities (PSD). B) Representative image from a 5xFAD heterozygous hippocampus displaying mitochondria with disrupted morphology and altered cristae organization. Insets in panels (A) and (B) illustrate examples of mitochondria categorized as healthy, moderately damaged, or severely damaged, based on cristae integrity scoring criteria. C) Quantification of mitochondrial cristae integrity in WT hippocampi revealed that >90% of mitochondria were classified as healthy, with few displaying moderate damage and none showing severe cristae disruption. Corresponding analysis of 5xFAD hippocampi demonstrated a significantly higher proportion of mitochondria exhibiting moderate disruption (p = 0.002) and fewer healthy mitochondria (p=0.139), compared to WT. Approximately 33% of all mitochondria in 5xFAD samples were moderately or severely damaged. Table indicates mean ± SD for percentages of mitochondria in each category of health / damage. D) 5xFAD mitochondria were significantly smaller in area than WT (p = .0448). * indicates p < .05 for difference between genotypes. Mean ± SD. N = 3-6.

Semi-quantitative evaluation of mitochondrial cristae integrity revealed significant differences between groups. In WT hippocampi, >90% of mitochondria were classified as healthy, with only a small proportion showing moderate or severe damage (Fig. 4C). In contrast, mitochondria from 5xFAD hippocampi exhibited significantly greater levels of cristae disruption, with approximately 45% categorized as moderately or severely damaged (Fig. 4D). Statistical analysis confirmed that the proportion of mitochondria that were healthy was significantly lower in 5xFAD mice than WT (Fig. 4C; t(5.259) = 3.622, p = .0139). The percentage of moderately damaged mitochondria was significantly higher in 5xFAD mice compared with WT controls (*t*(6.379) = 5.018, p = .002) and there was a trend for a higher proportion of severely damaged mitochondria (*t*(5) =1.7, p = .1499). Damaged mitochondria often exhibited shortened or absent cristae and vacuolated inner compartments. The 5xFAD mitochondria were significantly smaller in average area than WT (Fig. 4D; t(676.4) = 2.01, p = .0448).

Collectively, these findings demonstrate that synaptic mitochondria in young 5xFAD mice displayed substantial ultrastructural abnormalities indicative of early mitochondrial distress, preceding overt amyloid pathology.

### 3.5 There were regional differences in genotypic effects on dendritic complexity, length, and spine density at 1 month of age

There was a significant main effect of Genotype (p=.023) on intersections between concentric circles for Sholl analysis and pyramidal cell dendrites in CA1 (Fig. 5A), but not for DG granule cells (p=.063; Fig. 5D) or CA3 pyramidal cells (p=.31; Fig. 5G). This was also seen with area under the curve analyses of these interaction plots in CA1 (p=.034; Fig. 5B), DG (p=.077; Fig. 5E), and CA3 (p=.30; Fig. 5H). 5xFAD Het CA1 pyramidal cells had fewer intersections (Fig. 5A) and reduced area under the curve (Fig. 5B) than WT. All regions showed significant main effects of Distance from soma (CA1 (Fig. 5A); DG (Fig. 5D); CA3 (Fig. 5G); all p<.0001), with more interactions occurring closer to the soma. There was a significant reduction in dendritic length in 5xFAD Het DG granule cells (p=.014; Fig. 5F), as compared to WT. There were no differences between genotypes in dendritic length of pyramidal neurons in CA1 (p=.55; Fig. 5C) or CA3 (p=.70; Fig. 5I). There was a significant increase in synaptic density on 5xFAD Het apical dendrites in the DG (p=.02; Fig. 6B), as compared to WT. There were no genotypic differences in synaptic density in the CA1 (p=.56; Fig. 6A) and CA3 (p=.16; Fig. 6C) pyramidal cell apical dendrites.

**Figure 5.**
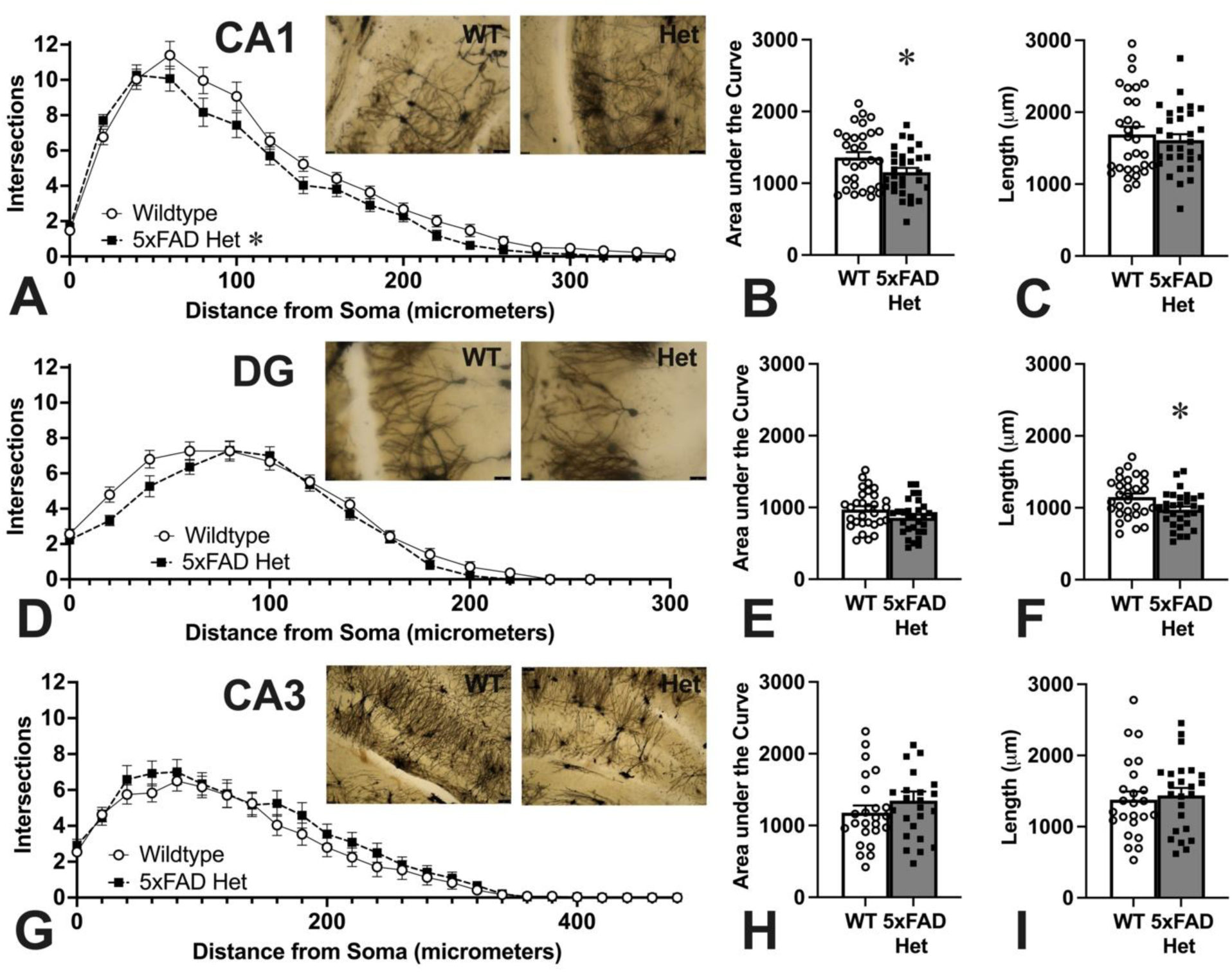
Sholl analysis yielded dendritic arborization (A,D,G), area under the curve from those arborization curves (B,E,H), and dendritic length measurements (C,F,I) for pyramidal neurons in CA1 (A,B,C) and CA3 (G,H,I) and granule cell neurons in DG (D,E,F) in 1 month old WT and 5xFAD Hets. Example Golgi-Cox stained images from WT and 5xFAD Het hippocampal regions are shown in insets within A,D,G (Bars=50µm). There were significantly fewer dendritic interactions with the concentric circles used in Sholl analysis (A) and area under the curve of all interactions (B) in 5xFAD Het, as compared to WT mice, in the CA1 region alone. DG granule cell dendrites showed a significantly shorter length in 5xFAD Hets than WT mice (F). N = 30 neurons (CA1 & DG), 24 neurons (CA3). Sholl analysis: Repeated measures ANOVA (CA1, CA3) or mixed effects ANOVA (DG). Area and length: unpaired t-test. Mean ± SEM.

**Figure 6.**
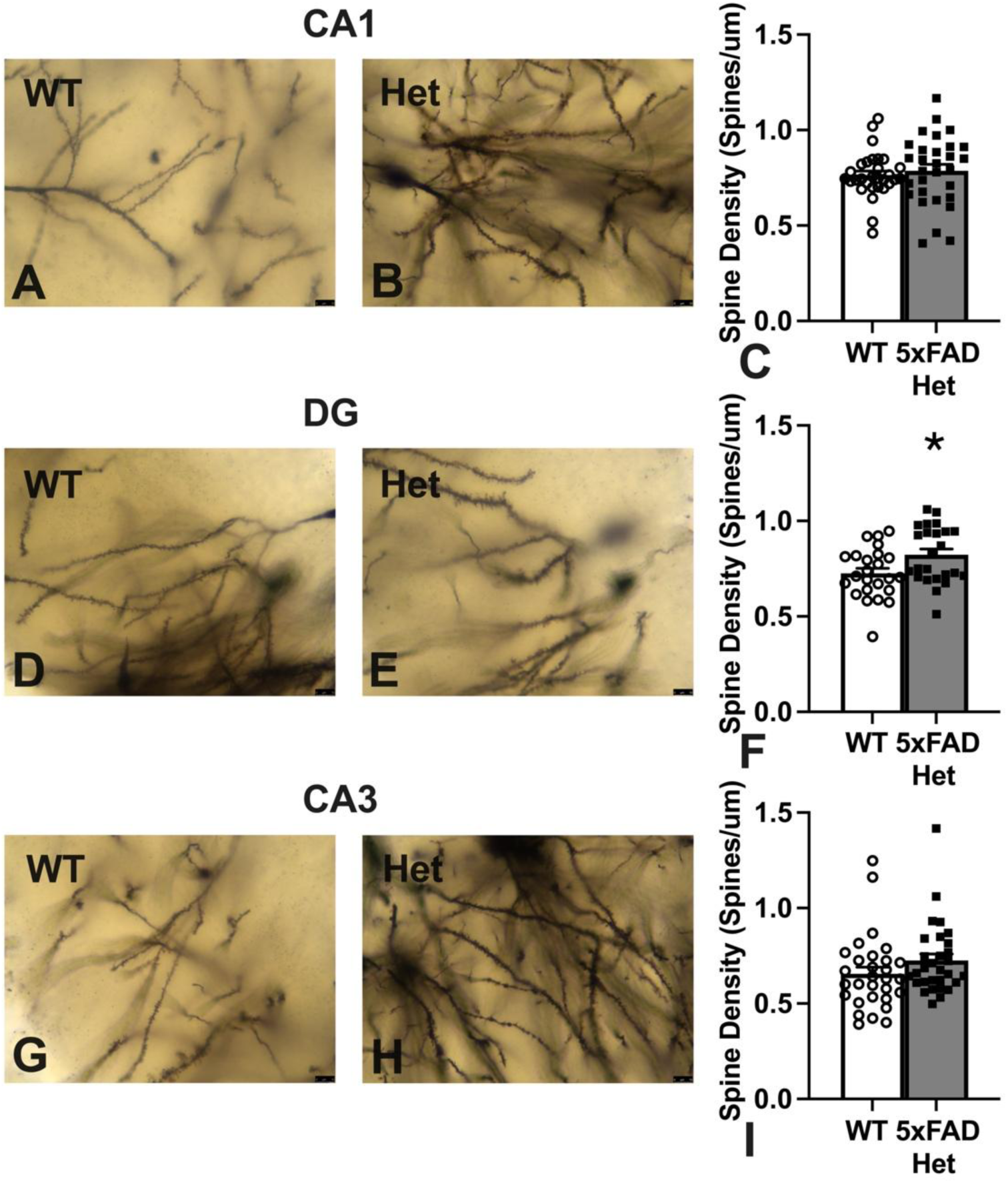
Spine density analyses from apical dendrites of CA1 pyramidal (A-C), CA3 pyramidal (G-I) and granule cell (D-F) neurons of the hippocampus from Golgi-Cox stained brains. Representative images from WT (A,D,G) and 5xFAD Hets (B,E,H) and graphs of spine density measures from CA1 (C), DG (F) and CA3 (I) regions. 5xFAD Hets had significantly higher density of spines in the DG than WT (F). Welch’s t-test. Mean ± SEM. N = 30 neurons (CA1 & CA3); 24 neurons (DG). Bars = 10 µm.

### 3.6 Spatial transcriptomics showed differential effects of genotype on hippocampal subregions at 1 month of age

*Global Data Visualization* - Global data was first grouped by similarity to determine if differences existed between subregions for further analysis. Both tSNE and UMAP (Fig. 7B,C) and hierarchical clustering (Fig. 7D) showed grouping along subregions, providing rationale for further investigation with differential gene expression analysis.

**Figure 7.**
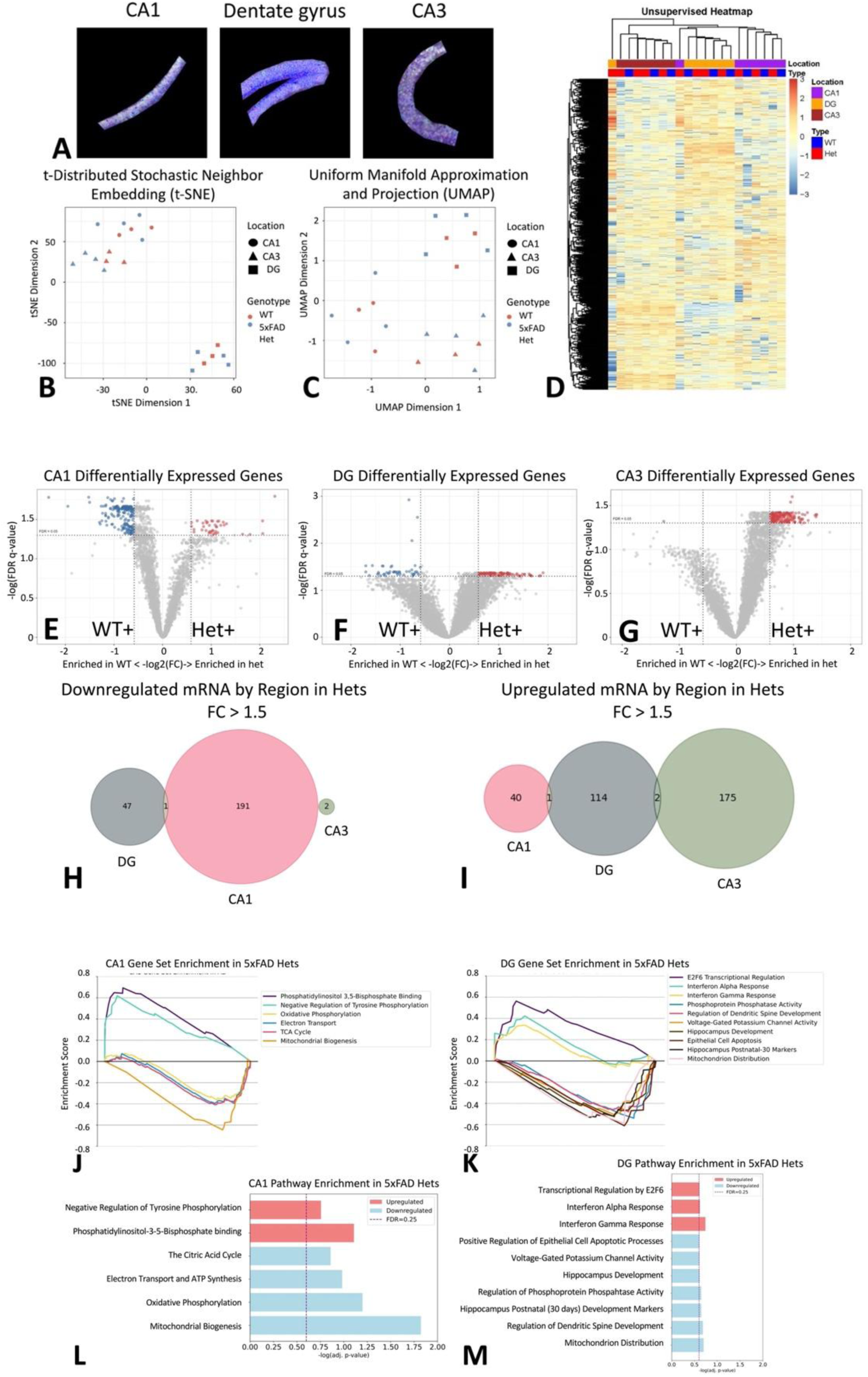
Spatial transcriptomics of subregions of the hippocampus. A) Representative images of regions of interest from CA1, dentate gyrus and CA3 subregions stained with NeuN, GFAP, Iba1 and DNA. B,C) Visualization of dimensionally reduced data using t-distributed stochastic neighbor embedding (t-SNE; B) and uniform manifold approximation and projection (UMAP; C) showed subregion separation, but not genotypic. D) Visualization of global log-fold changes separated by genotype and subregion, created using unsupervised hierarchical clustering, showed subregion differences. E-G) Volcano plots showing differential gene expression by subregion in the CA1 (E), DG (F), and CA3 (G). Double filtering was set to include |FC| > 1.5 and FDR q-value <0.05. Dots in red represent transcripts that are significantly increased in Hets with double filtering, and dots in blue represent transcripts that are significantly decreased in Hets with double filtering. H-I) Venn Diagrams showing number and overlap of differentially expressed genes within each subregion that had fold-changes greater than 1.5 with WT > 5xFAD Hets (H) or with 5xFAD Hets > WT (I) showed little overlap between subregions. Data visualized using Matplotlib and Matplotlib – Venn. J-M) Summary of Gene Set Enrichment Analysis Pathway Enrichments. Plots show the enrichment scores of pathway genes by location in the CA1 (J) and DG (K) subregions and the statistical significance of each pathway in the CA1(L) and DG(M) subregions. Gene sets with significant upregulation in 5xFAD Hets have positive enrichment scores, while significantly downregulated gene sets in 5xFAD Hets have negative enrichment scores (A-B). Data visualized using ggplot2, unless otherwise noted. N = 3-4.

#### Differential Gene Expression

Analyzing differentially expressed genes by genotype using NOISeqBio revealed many changes in transcript counts by subregion and genotype in 1 month old mice (Fig. 7E-M). Global expression and significance of transcripts were visualized in volcano plots (Figure 7E-G). In CA1, 191 transcripts were decreased in 5xFAD Hets vs WT and 40 were increased (Fig. 7E,H,I). In DG, 47 transcripts were decreased in 5xFAD Hets vs WT and 114 were increased (Fig. 7F,H,I). In CA3, 2 transcripts were decreased in 5xFAD Hets vs WT and 175 were increased (Fig. 7G,H,I). Differences in transcript counts between subregions had little overlap (Fig. 7H,I), so Gene Set Enrichment Analysis was performed to look at possible functional differences by subregions.

#### Gene Set Enrichment Analysis

GSEA revealed several differentially regulated pathways in CA1 (Fig. 7J,L) and DG (Fig. 7K,M), but none in CA3 (Data not shown) in 1 month old mice. The CA1 region showed the most significant effects of genotype on gene sets (Fig. 7L), compared to DG (Fig. 7M). The 5xFAD Hets exhibited higher mRNA levels than WT for genes involved in negative regulation of tyrosine phosphorylation and phosphatidylinositol-3-5-bisphosphate binding (Fig. 7L, Table 1S). The down-regulated gene sets in the 5xFAD Hets included citric acid cycle, electron transport and ATP synthesis, oxidative phosphorylation and mitochondrial biogenesis (Fig. 7L, Table 1S). Upregulated gene sets in the 5xFAD Het DG included interferon alpha and gamma response and downregulated mRNAs included 7 gene sets, including those involved in voltage-gated potassium channels, protein phosphatase activity regulation and mitochondrion distribution (Fig. 7M, Table 1S).

### 3.7 Single cell deconvolution suggests CA1 pyramidal neurons are the most vulnerable population to mitochondrial changes

To determine if the observed mitochondrial defects were ubiquitous or cell-type specific, we first reconstructed the cellular architecture of the hippocampus. Spatial deconvolution confirmed expected regional distributions, with *CA1.ProS* neurons, *CA3* neurons, and *DG* granule cells localizing to the greatest degree in their respective anatomical subregions (Fig. 8A). Importantly, we observed no significant shifts in the proportional abundance of these cell types between genotypes, indicating that the mutation does not drive overt neurodegeneration or cell loss at this timepoint.

**Figure 8.**
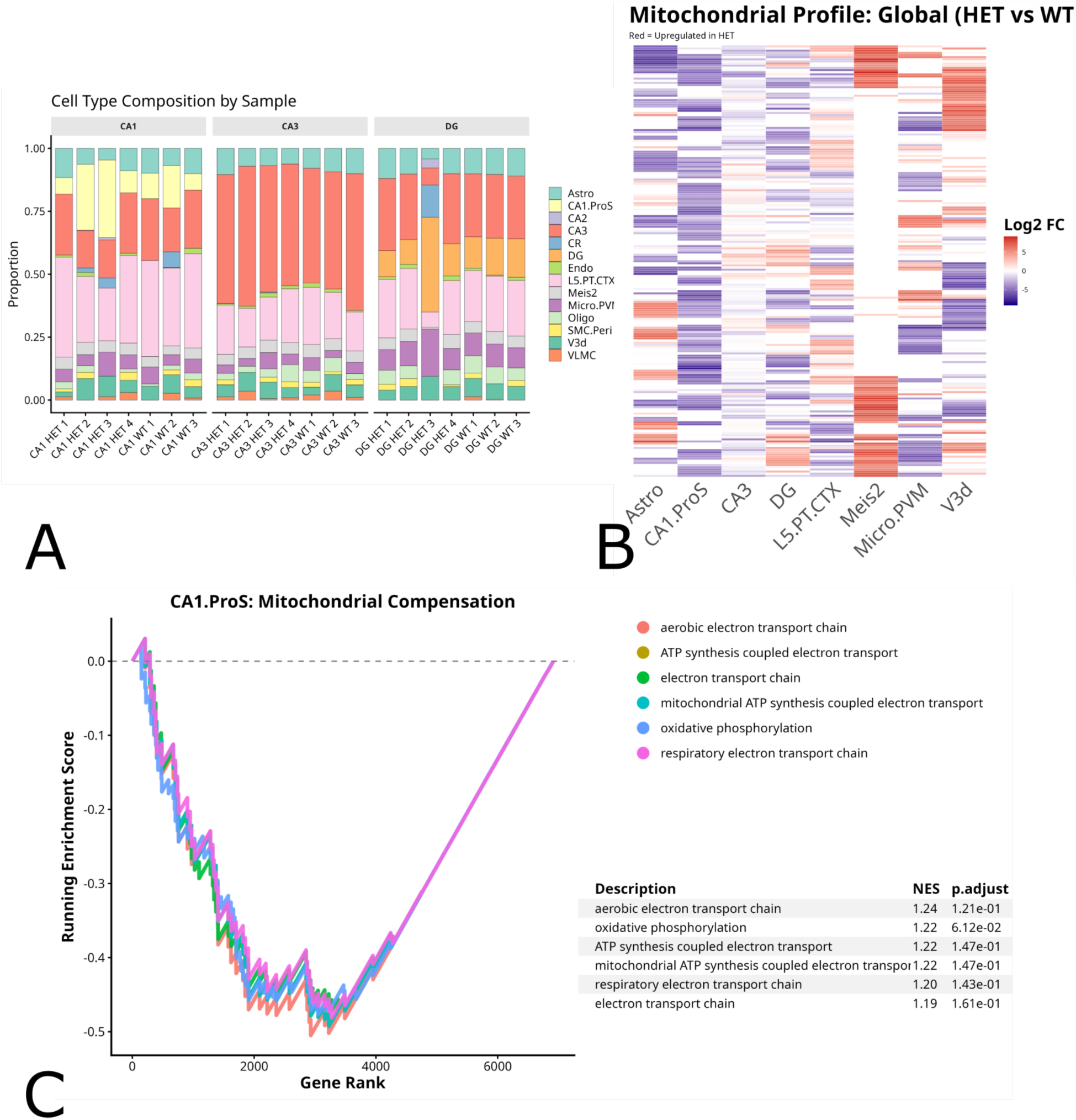
CA1 Pyramidal Neurons Exhibit a Unique, Compensatory Mitochondrial Downregulation. A) Cell type composition analysis of hippocampal subregions (DG, CA3, CA1) derived from spatial transcriptomic deconvolution. Stacked bar charts display the proportional abundance of major neuronal and glial populations across individual samples, confirming consistent cellular architecture between Wild-Type (WT) and Heterozygous (HET) genotypes. B) Heatmap of differential expression for a targeted panel of mitochondrial genes involved in transport, biogenesis, and oxidative phosphorylation. Columns represent cell types; rows represent individual genes. Color intensity indicates Log2 Fold Change (Log2FC) in HET relative to WT (Red = Upregulated in HET; Blue = Downregulated). While most cell types show neutral or heterogeneous patterns, CA1.ProS neurons display a coordinated downregulation across the majority of the panel. Statistical analysis confirms that CA1.ProS neurons are the primary driver of this variance (χ^2^ = 219.54, p < 0.0001), with a significantly higher proportion of upregulated transcripts compared to all other cell types (Fisher’s Exact Test, p < 0.0001). C) Gene Set Enrichment Analysis (GSEA) of the CA1.ProS expression profile. The running enrichment score (solid lines) indicates a strong positive enrichment for canonical mitochondrial pathways, including “Oxidative Phosphorylation” and “Aerobic Electron Transport Chain” in HET neurons. The table (right) displays normalized enrichment scores (NES) and adjusted p-values for significant pathways, confirming a robust compensatory metabolic response specifically within the CA1 pyramidal population. N = 3-4.

However, analysis of cell-type specific gene expression revealed a profound metabolic divergence. A heatmap of mitochondrial transcripts from the GSEA results in Figure 7 demonstrated that while most cell types displayed heterogeneous or neutral expression changes, *CA1.ProS* neurons exhibited a striking, coordinated downregulation across the majority of the panel (Fig. 8B). Statistical analysis confirmed this observation; the distribution of up- versus down-regulated mitochondrial genes was significantly dependent on cell type (χ^2^ = 219.54, p < 0.0001). Post-hoc residual analysis identified *CA1.ProS* neurons as the primary driver of this variance (Z = 10.79, p < 0.0001), exhibiting a significantly higher proportion of downregulated transcripts compared to the global tissue average. Fisher’s Exact test further confirmed that *CA1.ProS* neurons were significantly more likely to exhibit mitochondrial downregulation compared to all other cell types combined (p < 0.0001).

To define the functional consequence of this downregulation, we performed Gene Set Enrichment Analysis (GSEA) on the *CA1.ProS* expression profile. This analysis revealed a robust, negative enrichment for canonical energy production pathways, including “Aerobic Electron Transport Chain” (NES = 1.24) and “Oxidative Phosphorylation” (Fig. 8C). The specific induction of these pathways in HET CA1 neurons—but not in neighboring CA3 neurons or glia—suggests a localized compensatory mechanism where pyramidal neurons downregulate mitochondrial respiratory machinery in response to excitatory damage.

## 4 Discussion

Initially, research and subsequent treatments were focused on insoluble Aβ plaques as the initiating factor for AD (Krafft et al., 2022), but newer evidence suggests that these may be the body’s responses to injury, rather than the inciting causes (Mondragon-Rodriguez et al., 2010). Evidence is growing that Aβ42 or Aβ40 oligomers can trigger the neurodegeneration and cognitive deficits that are characteristic of AD (Tolar et al., 2021, Krafft et al., 2022, Haessler et al., 2024, Mroczko et al., 2018). In this study, we showed that the transgene was expressed as early as 15 days of age, but the accumulation of plaques did not become significant until 4 months of age in the 5xFAD Hets. Despite this lack of plaques, the 5xFAD Hets showed glutamate receptor hyperactivity, mitochondrial damage and associated RNA changes, and alterations in dendritic complexity at 1 month of age. In addition, we found regional differences within the hippocampus, with the CA1 region showing more changes in dendrites and RNA than CA3 or dentate gyrus at this early age. Given the pre-plaque age, these changes are likely to be related to Aβ oligomers.

### 4.1 Glutamate receptor hyperactivity

The 1 month old 5xFAD Het mice showed increased synaptic responses in both AMPA receptors and NMDA receptors expressing GluN2B subunits. There is evidence that this hyperactivity may be relevant to the development of AD, as humans with mild cognitive impairment, a preclinical stage of AD, show higher activation of brain regions while performing cognitive tasks than those with normal cognition (Dickerson et al., 2005, Foster et al., 2018, Mormino et al., 2012). PSEN mutant mice also show NMDA receptor hyperactivity by 3-5 months of age (Auffret et al., 2010, Auffret et al., 2009, Dewachter et al., 2008). Our results are thus in keeping with a prior study using PS1A246E PSEN mutant mice which also showed an increase in NMDA receptor GluN2B subunit expression (Dewachter et al., 2009). This is also associated with enhanced early LTP (1 hour post-stimulation), but reduced late LTP, suggesting that the hyperactivity is not beneficial for long-term synaptic plasticity (Auffret et al., 2010). Although NMDARs are normally good for initiating memories and synaptic plasticity (Morris et al., 1986, Alessandri et al., 1989, Butelman, 1989, Heale and Harley, 1990, Mondadori et al., 1989), overstimulation by glutamate can lead to calcium overload-related excitotoxicity and cell death (Choi, 1992). The 5xFAD mice express 2 different PSEN mutations which may explain the NMDA receptor hyperactivity in these mice (Oakley et al., 2006).

Aβ oligomers can increase NMDA and AMPA receptor-related calcium currents and membrane depolarization and calcium overload of mitochondria in organotypic cultures (Alberdi et al., 2010). There is also evidence that Aβ can induce reactive oxygen species via interaction with NMDA receptors (Shelat et al., 2008). Soluble Aβ can increase GluN2B subunit-related responses, which is linked to LTP inhibition (Li et al., 2011). This increase in response was seen in young 5xFAD Hets in the current study. Consequences of interactions between Aβ and GluN2B-containing receptors also include increasing mitochondrial calcium and membrane depolarization (Ferreira et al., 2015) and causing synaptic depression via non-ionotropic NMDA receptor functions (i.e., not via its ion channel) (Kessels et al., 2013). The fact that AMPA receptor activity was also elevated in young 5xFAD Hets, however, highlights a difference from PSEN mutants, which show no changes in AMPA receptors (Auffret et al., 2010, Auffret et al., 2009, Dewachter et al., 2008). Aβ has been shown to increase glutamate in the synaptic cleft by either enhancing astrocytic glutamate release (Talantova et al., 2013) or decreasing uptake via transporters (Li et al., 2011, Li et al., 2009). Excess synaptic glutamate may explain the increased responses in both AMPA and NMDA receptors in the 1 month old 5xFAD Hets. The NMDA receptor hyperactivity in PSEN mutants have been linked to the inability of the endoplasmic reticulum to buffer calcium (Costa et al., 2012, Schneider et al., 2001). However, it could also be due to deficits in mitochondrial calcium buffering (Stanika et al., 2009, Ma et al., 2020, Ferreira et al., 2015, Ryan et al., 2020, Mustaly-Kalimi et al., 2025, Walters and Usachev, 2023), particularly in the synaptic environment.

### 4.2 Mitochondrial injury and early bioenergetic compromise

Using a multi-modal strategy combining scanning transmission electron microscopy, spatial transcriptomics, and high-resolution neuronal morphometrics, our study shows early, subregion-specific hippocampal pathology in one-month-old 5xFAD heterozygous mice. Synaptic mitochondria display pronounced ultrastructural damage — including cristae disruption, vacuolization, and increased electron density — despite no net loss in mitochondria number per image field (Fig. 4). Spatial transcriptomics reveals distinct, subfield-dependent transcriptional responses: CA1 is marked by pronounced downregulation of mitochondrial and bioenergetic gene programs; DG shows upregulation of interferon-related immune pathways along with dysregulation of mitochondrial distribution and ion channel genes; while the CA3 region remains largely unaffected. Morphologically, CA1 pyramidal neurons exhibit reduced dendritic complexity, DG granule cells show reduced dendritic length but increased apical dendritic spine density, and CA3 neurons are comparatively spared. Together, these findings indicate that synaptic mitochondrial distress, metabolic reprogramming, and local immune activation combine early — well before overt amyloid plaque pathology — to drive structural synaptic remodeling in the 5xFAD hippocampus.

The striking ultrastructural abnormalities seen in 5xFAD synaptic mitochondria provide powerful evidence that mitochondrial integrity is compromised at the very earliest stages of pathology. Cristae integrity is critical for efficient electron transport chain function and ATP production; loss or disruption of cristae, vacuolization, and membrane density changes likely impair oxidative phosphorylation (OXPHOS) and reduce local ATP availability at synapses. Concurrently, our spatial transcriptomic data show significant downregulation in CA1 of gene sets related to the citric acid cycle, electron transport, ATP synthesis, and mitochondrial biogenesis (Fig. 7L), suggesting both a functional and transcriptional suppression of the mitochondrial energy apparatus. By employing spatial deconvolution, we further resolved this transcriptomic suppression to the cellular level (Fig. 8). We found that this metabolic downregulation is not a pan-cellular phenomenon but is specifically driven by CA1 pyramidal neurons (CA1.ProS). While neighboring CA3 neurons and glial populations maintained relatively stable mitochondrial gene profiles, CA1 pyramidal neurons exhibited a coordinated repression of oxidative phosphorylation and electron transport chain transcripts. The fact that these alterations are intrinsic to CA1 pyramidal neurons and restricted to synaptic mitochondria (without reduced mitochondrial number) argues strongly that energy failure at the level of CA1 pyramidal synapses — rather than global neuronal mitochondrial depletion — is one of the earliest detectable pathologies in the 5xFAD hippocampus.

These observations align with prior work in AD models. Studies using synaptosomes from 4 and 9 month old 5xFAD mice brains demonstrated early mitochondrial dysfunction, including reduced respiration and impaired synaptic mitochondrial bioenergetics, that deteriorates with increased age (Wang et al., 2016). In human AD tissue, alterations in mitochondrial–ER contacts and reorganized mitochondria have also been documented, suggesting structural mitochondrial changes are a conserved feature of early disease (Area-Gomez et al., 2012). More broadly, mitochondrial dysfunction and structural abnormalities — including cristae loss — have long been implicated as central contributors to neurodegenerative disease pathogenesis, by promoting energy failure and enhancing reactive oxygen species (ROS) production (Rhein et al., 2009, Porcellotti et al., 2015, Ikon and Ryan, 2017, Cole et al., 2010).

### 4.3 Possible mechanisms underlying cristae destruction and mitochondrial ultrastructural damage

In addition to early metabolic suppression, our morphometric analysis revealed striking disruption of mitochondrial cristae in synaptic compartments. The literature identifies several converging mechanisms that could account for this ultrastructural damage. First, alterations in mitochondrial dynamics or cristae-organizing complexes can destabilize inner membrane architecture, producing the fragmented or vacuolated cristae seen in neurodegenerative models (Li et al., 2004, Porcellotti et al., 2015). Second, impaired mitophagy may allow structurally compromised mitochondria to accumulate, especially under conditions of high synaptic energetic demand (Du et al., 2008, Zhang et al., 2016). Third, Aβ itself can directly perturb mitochondrial membrane curvature and fluidity, reducing the stability of cristae-like invaginations even before overt plaque deposition (Rodrigues et al., 2001, Khalifat et al., 2012). Additionally, subtle disruptions in calcium handling may promote opening of the mitochondrial permeability transition pore, leading to osmotic swelling and loss of cristae structure (Rhein et al., 2009). Finally, oxidative stress—exacerbated by impaired electron transport—can damage inner membrane lipids and proteins, accelerating cristae collapse (Rhein et al., 2009, Ikon and Ryan, 2017, Cole et al., 2010). Taken together, these mechanisms provide a plausible multifactorial explanation for the early cristae destruction observed in 5xFAD synaptic mitochondria and support the interpretation that synaptic bioenergetic failure is a primary, not secondary, feature of early pathology.

4.4 **Spatially heterogeneous inflammatory and metabolic transcriptomic programs**

Our spatial transcriptomics data reveal strong subfield specificity in early genotype-associated transcriptional changes. In CA1, the dominant signature is suppression of mitochondrial/energetic pathways; in DG, upregulation of interferon-related immune pathways occurs alongside dysregulation of mitochondrial distribution and ion channel–related gene sets (Fig. 7). The DG interferon signature is especially noteworthy given recent studies implicating type I/II interferon signaling in driving synapse loss, microglial activation, and cognitive impairment in amyloid-bearing models (Rhein et al., 2009, Minter et al., 2016, Roy et al., 2022, Roy et al., 2020). That DG — but not CA3 — shows this signature suggests differential subfield vulnerability and immune engagement at very early stages.

This dual pattern — metabolic suppression in CA1 and immune activation in DG — supports a model in which **bioenergetic failure at synapses** (particularly in CA1) co-occurs with **innate immune activation** (notably in DG), potentially acting together to destabilize synaptic structure. The spatial segregation suggests that different hippocampal subfields may engage either energetic compensation or immune-driven remodeling depending on local context (metabolic load, synaptic activity, Aβ exposure, calcium handling, microglial proximity). We also have evidence of genotype-dependent changes in multiple lipid species within individual hippocampal subregions, including CA1, consistent with early metabolic and membrane remodeling (Supplemental Fig. 1).

### 4.5 Dendritic changes

There were reductions in dendritic complexity in the CA1 regions and dendritic length in the dentate gyrus at 1 month of age in 5xFAD Hets. Declines in dendritic extent and complexity are characteristic of AD (Petrides et al., 2016, Coleman and Flood, 1987). Overexpression of Aβ in rodent models is also associated with decreased dendritic complexity (Golovyashkina et al., 2015, Wu et al., 2010). The differential susceptibility of CA1 neurons to dendritic simplification has been seen in mutant APP organotypic hippocampal slice cultures exposed to human tau (Golovyashkina et al., 2015). These dendritic changes could also be related to the glutamate receptor hyperactivity seen in the young 5xFAD Hets. Stress-induced dendritic retraction in rats is driven by NMDA receptor activation (Martin and Wellman, 2011). The dendritic simplification noted above was almost entirely prevented by NMDA receptor antagonists (Golovyashkina et al., 2015). Non-ionotropic NMDA receptor signaling, which can be triggered by Aβ (Kessels et al., 2013), leads to spine shrinkage via pathways including nitric oxide synthase 1 adaptor protein (NOS1AP), neuronal nitric oxide synthase (nNOS), nNOS enzymatic activity, and cofilin (Stein et al., 2020).

Aβ applied to hippocampal or Cos7 cell cultures induces spine elimination (Kawaguchi et al., 2022, Martinez et al., 2024). There is also evidence that Aβ effects via NMDA receptors can lead to dendritic spine loss (Prikhodko et al., 2024). This is contrary to our finding that dentate gyrus granule cells in the 1 month old 5xFAD Hets showed an increased spine density. There is evidence that acute application of Aβ to hippocampal organotypic cultures induces an increase in spine density in the CA1 neurons, involving a decrease in stable spines and an increase in dynamic spines, but they also document an increase in dendritic length (Ortiz-Sanz et al., 2020). It’s not clear whether the 5xFAD Hets had an actual increase in spine numbers or whether it was related to the reduction in dendritic length in the dentate gyrus.

### 4.6 CA1 selective vulnerability

In both loss of dendritic complexity and changes in transcripts, the CA1 region in the 5xFAD Hets was more affected than CA3 or dentate gyrus neurons. The CA1 region is more vulnerable than these other hippocampal regions to inflammation, hypoglycemia, ischemia and excitotoxicity (Lahtinen et al., 2001, Kristensen et al., 2001, Bramlett et al., 1999, Alkadhi, 2019). CA1 pathology, atrophy, and synaptic and neuronal loss occur early in the development of AD (Scheff et al., 2007, Braak and Braak, 1997, Kerchner et al., 2012, Kerchner et al., 2010, West et al., 1994). Our deconvolution analysis provides a mechanistic transcriptomic basis for this historical observation of selective vulnerability. We demonstrated that at one month of age, CA1 pyramidal neurons—but not CA3 neurons or DG granule cells—initiate a profound downregulation of mitochondrial respiratory machinery (Fig. 8B, C). It is plausible that the specific mitochondrial downregulation observed in CA1 pyramidal neurons represents a compensatory response to this excitotoxic drive—an attempt to mitigate ROS production in the face of calcium overload—or a direct failure of bioenergetics due to that overload. The selective susceptibility of CA1 to ischemia is associated with a lower expression of glutamate transporters compared to CA3 and DG regions (Zhang et al., 2011). There are also more GluN2B-containing NMDA receptors in CA1 than in dentate gyrus (Coultrap et al., 2005). The dendritic simplification in the CA1 region of the young 5xFAD Hets may thus be related to the hyperactivity in glutamate receptors at this age. Calbindin, a neuronal calcium-binding protein, is found in almost all DG granule cells, but only in a third of CA1 pyramidal cells (Sloviter, 1989) and the granule cells recover faster than CA1 pyramidal cells following depolarization-related calcium elevations (Baba et al., 2002), suggesting that calcium buffering capacity is reduced in CA1 neurons, compared to DG. This could be relevant to both the glutamate receptor hyperactivity and mitochondrial damage seen in the one-month-old 5xFAD Hets.

### 4.7 Linking mitochondrial dysfunction to neuronal and synaptic structural remodeling

The structural phenotypes we observe — reduced dendritic complexity in CA1, truncated dendrites but increased apical spinogenesis in DG, and relative sparing of CA3 — align well with the molecular and ultrastructural signatures. In CA1, energy deficiency from impaired mitochondrial function likely limits the maintenance of distal dendritic arbors and stable synapses. In DG, the combination of metabolic stress and immune activation may trigger synaptic reorganization, possibly forming compensatory but less stable synapses (manifested as increased spine density), or alternatively reflecting dysregulated synaptogenesis driven by inflammatory cues. The relative preservation of CA3 supports the idea of region-specific resilience — perhaps owing to lower metabolic demand, different calcium handling, less Aβ burden, or more robust quality-control capacity in that subfield. Importantly, such heterogeneity may underlie the well-documented differential vulnerability of hippocampal subfields in human Alzheimer’s disease, and suggests that early pathology may unfold along spatially selective trajectories.

### 4.8 Mechanistic implications and therapeutic insight

Our data support a model in which synapse-localized mitochondrial injury triggers a cascade of events: bioenergetic failure, suppressed mitochondrial gene expression, altered synaptic transmission, and region-specific innate immune activation. This convergence likely destabilizes synaptic architecture and may set the stage for later, irreversible neurodegeneration. From a therapeutic standpoint, this suggests that preservation of mitochondrial integrity at synapses — via interventions that support mitochondrial dynamics, promote mitophagy, stabilize inner membranes, or buffer calcium and oxidative stress — could be especially effective if applied early. Additionally, immune modulation, particularly inhibiting pathologic interferon signaling in vulnerable regions like DG, might prevent maladaptive synaptic remodeling.

### 4.9 Impacts on the animal

It is unclear how these changes to the hippocampus impact the animal as a whole. There is one group that reports spatial memory deficits as early as 1 month of age in the 5xFAD Het mice (Tang et al., 2016), although we have not been able to detect these declines at 2.5 months (KRM, personal observation). We recently reported that both male and female 5xFAD Hets show increases in sleep fragmentation prior to Aβ plaque deposition at 1 month of age (Kim et al., 2025). Sleep disruption appears to be an early sign of AD (Khosroazad et al., 2023, Berg, 2008, Lim et al., 2013) and decreased sleep duration is related to Aβ buildup within the hippocampus and other brain regions (Dufort-Gervais et al., 2019, Winer et al., 2021, Shokri-Kojori et al., 2018, Spira et al., 2013). It is not clear whether Aβ is the initiating cause of the sleep fragmentation, however, because sleep disruptions interfere with the glymphatic system, which is involved in clearance of Aβ in the brain during rest periods (Cordone et al., 2019, Xie et al., 2013). We have shown that the 5xFAD Hets on the C57BL/6J background expressed the transgene by 15 days of age and others show intraneuronal Aβ detectable at 1 month of age (Caccavano et al., 2020). Thus it is possible that early Aβ production contributes to sleep disturbances, which then exacerbates the accumulation of Aβ (Insel et al., 2021, Harenbrock et al., 2023).

There are potential relationships between sleep fragmentation and the other abnormalities seen at 1 month of age in the 5xFAD Hets. Chronic sleep deprivation can lead to dendritic simplification (Brodin et al., 2025). NMDA receptors in the hypothalamus are essential for normal sleep/wake cycling (Miracca et al., 2022) and systemic application of an NMDA receptor modulator facilitates sleep (Burgdorf et al., 2019). However, sleep deprivation can lead to AMPA receptor hyperactivity in the cerebral cortex (Vogt et al., 2025) and chronic insomnia is associated with increased NMDA receptor GluN1 subunit and glutamate in peripheral blood (Lin et al., 2024). Individuals with primary mitochondrial dysfunction frequently experience breathing issues that fragment sleep (Brunetti et al., 2021); however, induced sleep deprivation is also associated with mitochondrial morphology changes in the cortex and hippocampus (Lu et al., 2021, Aboufares El Alaoui et al., 2023, Sarnataro, 2025). It remains to be determined whether these changes are linked and, if so, which comes first.

### 4.10 Limitations

Several important caveats should be acknowledged. First, although ultrastructural and transcriptomic data strongly suggest impaired oxidative phosphorylation, we did not directly measure synaptic ATP levels, mitochondrial membrane potential, or ROS production in situ. Without functional assays (e.g., synaptosomal respirometry, genetically encoded ATP/ROS sensors, or redox imaging), the link between cristae disruption and energetic failure remains inferential. Second, although we propose multiple mechanisms for cristae destruction, receptor hyperactivity, and dendritic simplification, we cannot yet distinguish among them in our model; detailed mechanistic experiments will be needed to test their relative contributions. Finally, these results involve combined sexes. Our sleep/wake study shows sex differences in the severity of sleep disruption by 4 and 6 months of age in the 5xFAD Hets. Future studies should focus on each sex individually.

## 5 Conclusion

In summary, our integrated synaptic activity, structural, transcriptomic, and morphometric analyses reveal that glutamate receptor hyperactivity, synaptic mitochondrial damage, suppressed bioenergetic gene programs, and region-specific immune activation are among the earliest detectable perturbations in the 5xFAD hippocampus — occurring well before overt amyloid pathology or neuronal loss. These observations highlight synaptic mitochondria and innate immune signaling as convergent early contributors to synaptic instability in amyloid-driven neurodegeneration. Targeting mitochondrial preservation and immune modulation may therefore represent promising strategies for early intervention in Alzheimer’s disease.

## Supplemental Material

Pilot MALDI mass spectrometry imaging (MSI) reveals early lipidomic remodeling in the pre-plaque 5xFAD hippocampus. Using high-resolution MALDI-MSI on snap-frozen brain sections from 1-month-old 5xFAD and WT mice, hippocampal subregions (CA1, CA3, DG) were spatially resolved and segmented, enabling subregion-specific lipid profiling. Multivariate analysis demonstrated clear separation of 5xFAD versus WT lipid signatures, as well as separation among hippocampal subregions, indicating that lipid composition is altered early and in a region-dependent manner.

Unsupervised clustering and differential analysis identified genotype-dependent changes in multiple lipid species within individual hippocampal subregions, including CA1, consistent with early metabolic and membrane remodeling. Although exploratory (N=1), these data provide independent spatial evidence that molecular alterations extend beyond transcripts to lipid metabolism at a pre-plaque stage, reinforcing the concept that CA1 neurons experience early bioenergetic and membrane stress. When integrated with spatial transcriptomics showing downregulation of oxidative phosphorylation, TCA cycle, electron transport, and mitochondrial biogenesis pathways in CA1, the lipid MSI data support a multi-omic convergence on early mitochondrial and metabolic dysfunction localized to CA1.

## Acknowledgements

The authors would like to thank Drs. Cedric Boluda and Segolene Chaussis for their excellent technical assistance with the study. NIH NIA (5R21AG060206-02 to T.M.H.) supported EM and Golgi stain experiments. Oregon State University Department of Biochemistry & Biophysics provided undergraduate and graduate student support. Layton Alzheimer’s Disease Center (E.S.) and Oregon Partnership for Alzheimer’s Research (K.R.M) supported electrophysiological studies. Oregon State University Biomedical Sciences supported veterinary and graduate students. Linus Pauling Institute at Oregon State University provided funding for spatial transcriptomic experiment. NIH HEI grant (S10OD032323 to C.S.M.) and resources in OSU’s mass spectrometry center supported in part by institutional funds supported the spatial metabolomics experiment.

---

**Supplemental Figure 1.**
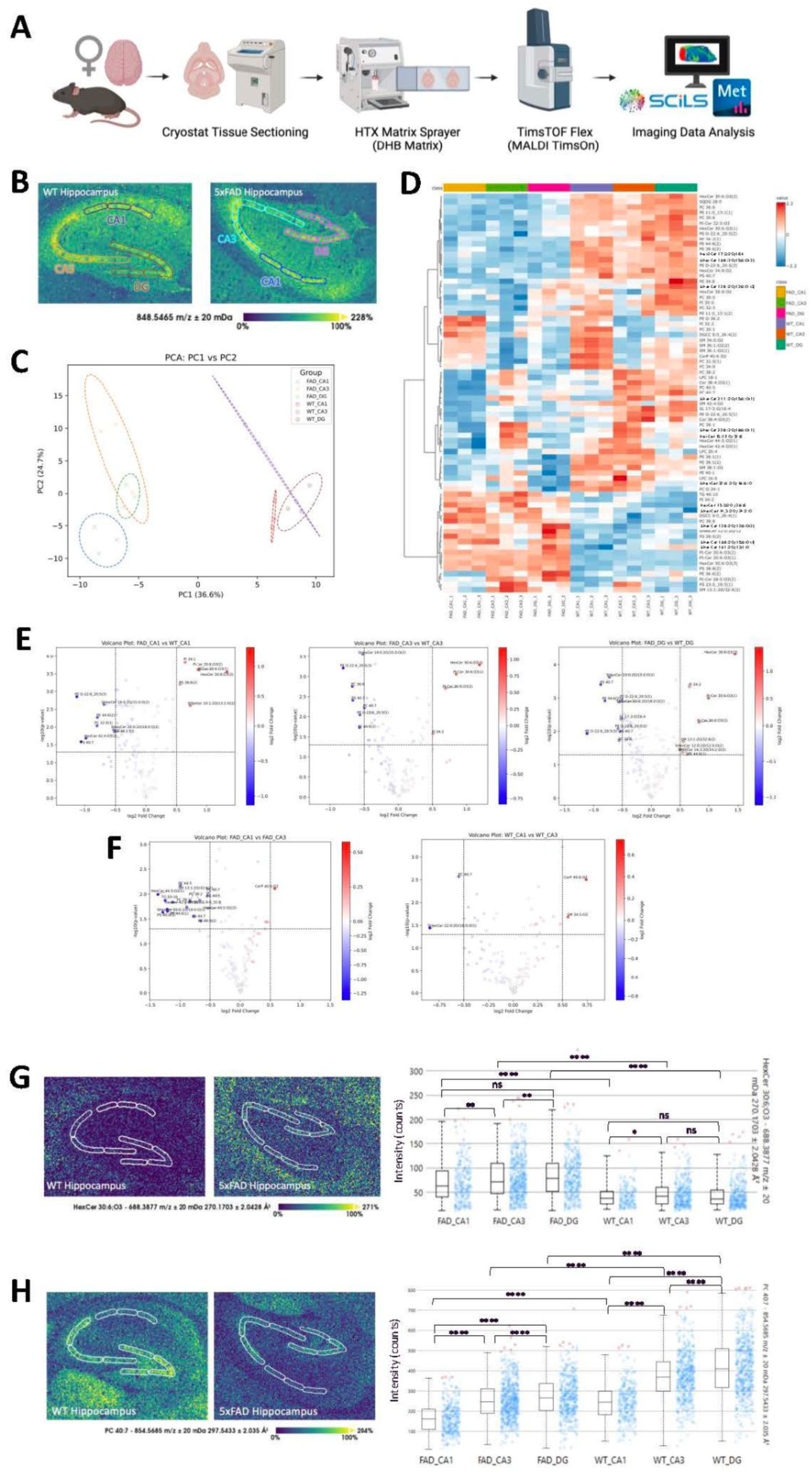
Mass spectrometry imaging data illustrates lipid differences within hippocampal regions in 1 month old 5xFAD mouse. (A) MALDI-MSI (matrix assisted laser desorption/ionization – mass spectrometry imaging) workflow for brain tissue analysis (created with the BioRender scientific illustration software). Female snap frozen brain tissue of 1 month old 5xFAD mouse and WT counterpart were sectioned horizontally into 12 μm-thick slices at -23 °C using a cryostat (Leica CM3050 S). Horizontal sections exposing the hippocampus region of interest were adhered to ITO slides (Bruker MALDI IntelliSlides), with comparative 5xFAD and WT sections on one slide. One slide (N=1) was collected, dried in a vacuum desiccator for 30 min, then stored at -80 °C until use. Before analysis, tissues were thawed in a vacuum desiccator for 15 min at room temperature. DHB matrix (40 mg/mL) was prepared in 70% MeOH in water and deposited onto the slide using the HTX M3+ Sprayer. Spraying parameters were 75 °C temperature, 10 psi pressure, 100 μL/min flowrate, 1200 mm/min velocity, 10 second drying time, and 8 passes. MSI data was collected on the TimsTOF flex mass spectrometer (Bruker Scientific, LLC, Bremen, Germany). The laser was set to 20 μm diameter and collected over a mass range of 100-1300 Da under positive ionization mode. The raw imaging data was processed in Bruker’s SCiLS Lab software for analysis and root mean square (RMS) normalization. Lipid MSI peaks were selected using 75% T-ReX^3^ feature finding and exported for annotation in Metaboscape with lipid species and MS-DIAL spectral library^1^ based on exact mass and collision cross section (CCS). (B) MSI data in SCiLS Lab resolved hippocampal subregions at m/z 849.5527. Based on this ion image, the CA1, CA3, and DG regions of WT and 5xFAD were segmented into three smaller subregions. The RMS normalized, average peak area intensity for all 18 subregions were generated and exported out of SCiLS for data visualization with Python. Prior to Python, the data was normalized using MetaboAnalyst 6.0 (low-abundance filtering by 10% of mean intensity value, normalization by sum, and log2 data transform). (C) Principal component analysis (PCA) plot showed separation of 5xFAD and WT hippocampus lipid profiles along PC1 and separation of hippocampal subregions along PC2. Hippocampal subregion separated along PC2 similarly between genotypes. (D) Heatmap of lipids, generated in MetaboAnalyst after ANOVA, show top 75 significantly expressed lipids between genotypes with hierarchical clustering of lipid features (Euclidean distance similarity measure and Ward’s linkage). (E) Volcano plots showing differentially expressed lipids between subregions (eg. FAD_CA1 vs WT_CA1). Double filtering was set to include |FC| > 0.5 and p-value < 0.05. Labeled dots in red represent lipids increased in Hets after double filtering, and labeled dots in blue represent lipids decreased in 5xFAD. (F) Volcano plots showing differentially expressed lipids between same genotype subregions. Labeled dots in red represent lipids increased in CA1 after double filtering, and labeled dots in blue represent lipids decreased in CA1. (G) From the volcano plots, a shared upregulated lipid in 5xFAD hippocampal subregions was identified as HexCer 30:6;O3 [M+K]+. The ion image validates external data analysis results, visualizing a higher intensity signal in the 5xFAD subregions. The corresponding ion intensity box plots generated in SCiLS Lab confirmed higher abundance of HexCer 30:6;O3 [M+K]+ in 5xFAD subregions. (H) Across both genotypes, a shared downregulated lipid in the CA1 subregion compared to the CA3 subregion was identified as PC 40:7 [M+Na]+. The corresponding ion intensity box plots generated in SCiLS Lab further confirmed lower abundance of PC 40:7 [M+Na]+ in the CA1 subregion across genotypes, but also an overall lower abundance in 5xFAD hippocampus. P-values were assessed in SciLS Lab using the two-sided t test: ns ≥ 0.05, * P < 0.05, ** P < 0.01, *** P < 0.001, **** P< 0.0001. (Tsugawa et al., 2015).

**Supplemental Table 1.**
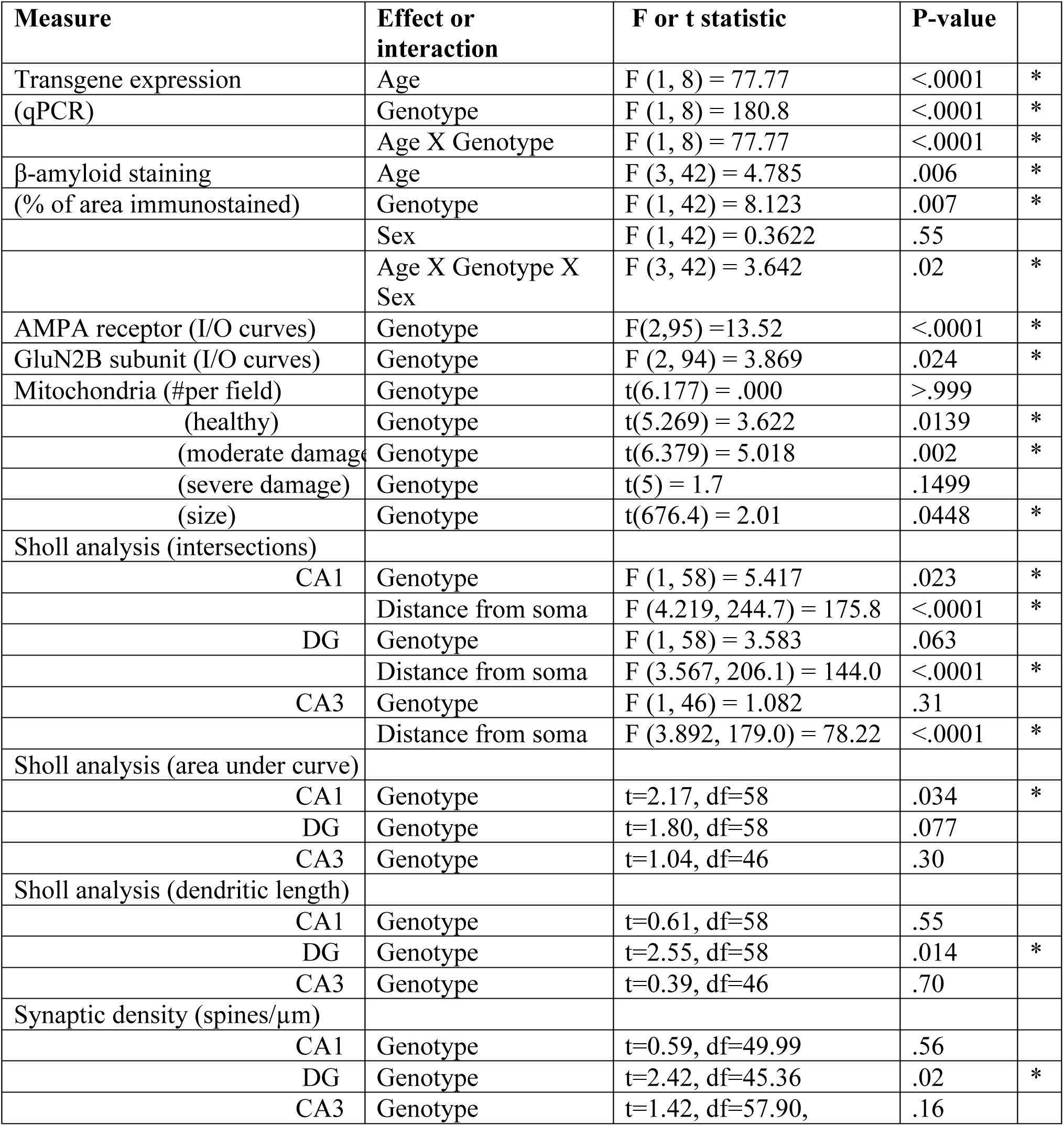
Statistical details.

**Supplemental Table S2:**
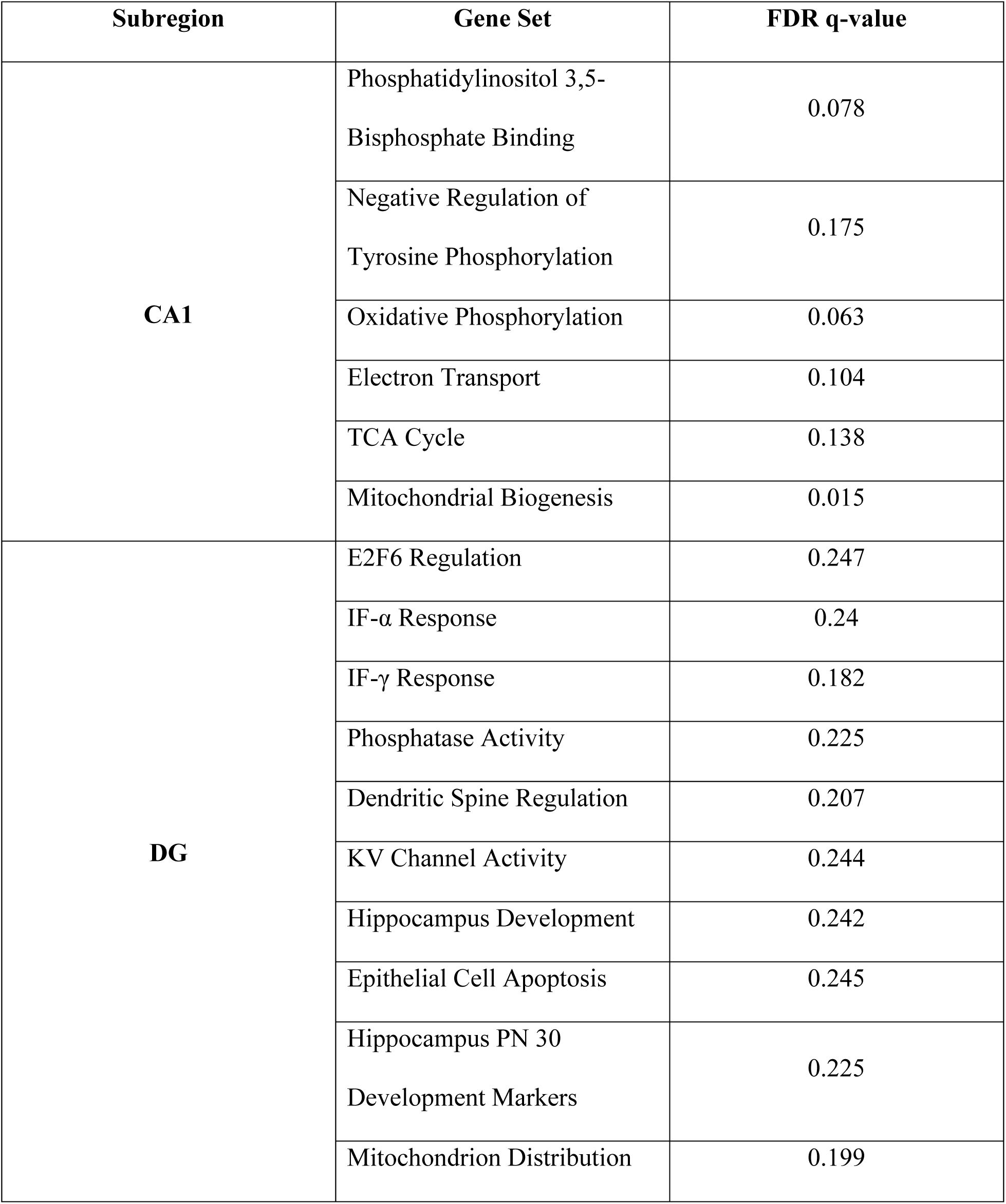
FDR Q-values associated with each GSEA pathway.

## Notes

https://metaspace2020.org/datasets?q=2026-02-03_18h41m55s

https://doi.org/10.7267/5999nc87c

### Competing Interest Statement

The authors have declared no competing interest.

