## Supplemental Materials for "Appearance of Amyloid-β Early in Life Initiates Neuronal Hyper-excitability, Mitochondrial Decay, and Loss of Dendritic Complexity in the Hippocampal CA1 Region of 5xFAD Mice"

### *Supplementary Material*

**Supplemental Table 1.** Statistical details

| Measure | Effect or interaction | F or t statistic | P-value | |
|---|---|---|---|---|
| Transgene expression | Age | F (1, 8) = 77.77 | <.0001 | * |
| (qPCR) | Genotype | F (1, 8) = 180.8 | <.0001 | * |
| | Age X Genotype | F (1, 8) = 77.77 | <.0001 | * |
| β-amyloid staining | Age | F (3, 42) = 4.785 | .006 | * |
| (% of area immunostained) | Genotype | F (1, 42) = 8.123 | .007 | * |
| | Sex | F (1, 42) = 0.3622 | .55 | |
| | Age X Genotype X Sex | F (3, 42) = 3.642 | .02 | * |
| AMPA receptor (I/O curves) | Genotype | F(2,95) =13.52 | <.0001 | * |
| GluN2B subunit (I/O curves) | Genotype | F (2, 94) = 3.869 | .024 | * |
| Mitochondria (#per field) | Genotype | t(6.177) = .000 | >.999 | |
| (healthy) | Genotype | t(5.269) = 3.622 | .0139 | * |
| (moderate damage) | Genotype | t(6.379) = 5.018 | .002 | * |
| (severe damage) | Genotype | t(5) = 1.7 | .1499 | |
| (size) | Genotype | t(676.4) = 2.01 | .0448 | * |
| Sholl analysis (intersections) | | | | |

| | | | | |
|---|---|---|---|---|
| CA1 | Genotype | F (1, 58) = 5.417 | .023 | * |
| | Distance from soma | F (4.219, 244.7) = 175.8 | <.0001 | * |
| DG | Genotype | F (1, 58) = 3.583 | .063 | |
| | Distance from soma | F (3.567, 206.1) = 144.0 | <.0001 | * |
| CA3 | Genotype | F (1, 46) = 1.082 | .31 | |
| | Distance from soma | F (3.892, 179.0) = 78.22 | <.0001 | * |
| Sholl analysis (area under curve) | | | | |
| CA1 | Genotype | t=2.17, df=58 | .034 | * |
| DG | Genotype | t=1.80, df=58 | .077 | |
| CA3 | Genotype | t=1.04, df=46 | .30 | |
| Sholl analysis (dendritic length) | | | | |
| CA1 | Genotype | t=0.61, df=58 | .55 | |
| DG | Genotype | t=2.55, df=58 | .014 | * |
| CA3 | Genotype | t=0.39, df=46 | .70 | |
| Synaptic density (spines/µm) | | | | |
| CA1 | Genotype | t=0.59, df=49.99 | .56 | |
| DG | Genotype | t=2.42, df=45.36 | .02 | * |
| CA3 | Genotype | t=1.42, df=57.90, | .16 | |

**Supplemental Table S2**: FDR Q-values associated with each GSEA pathway

| Subregion | Gene Set | FDR q-value |
|---|---|---|
| CA1 | Phosphatidylinositol 3,5-Bisphosphate Binding | 0.078 |
| | Negative Regulation of Tyrosine Phosphorylation | 0.175 |
| | Oxidative Phosphorylation | 0.063 |
| | Electron Transport | 0.104 |
| | TCA Cycle | 0.138 |
| | Mitochondrial Biogenesis | 0.015 |
| | E2F6 Regulation | 0.247 |
| | IF-α Response | 0.24 |
| | IF-γ Response | 0.182 |
| | Phosphatase Activity | 0.225 |

| | | |
|---|---|---|
| **DG** | Dendritic Spine Regulation | 0.207 |
| | KV Channel Activity | 0.244 |
| | Hippocampus Development | 0.242 |
| | Epithelial Cell Apoptosis | 0.245 |
| | Hippocampus PN 30 Development Markers | 0.225 |
| | Mitochondrion Distribution | 0.199 |

**Supplemental Material**

Pilot MALDI mass spectrometry imaging (MSI) reveals early lipidomic remodeling in the pre-plaque 5xFAD hippocampus. Using high-resolution MALDI-MSI on snap-frozen brain sections from 1-month-old 5xFAD and WT mice, hippocampal subregions (CA1, CA3, DG) were spatially resolved and segmented, enabling subregion-specific lipid profiling. Multivariate analysis demonstrated clear separation of 5xFAD versus WT lipid signatures, as well as separation among hippocampal subregions, indicating that lipid composition is altered early and in a region-dependent manner.

Unsupervised clustering and differential analysis identified genotype-dependent changes in multiple lipid species within individual hippocampal subregions, including CA1, consistent with early metabolic and membrane remodeling. Although exploratory (N=1), these data provide independent spatial evidence that molecular alterations extend beyond transcripts to lipid metabolism at a pre-plaque stage, reinforcing the concept that CA1 neurons experience early bioenergetic and membrane stress. When integrated with spatial transcriptomics showing downregulation of oxidative phosphorylation, TCA cycle, electron transport, and mitochondrial biogenesis pathways in CA1, the lipid MSI data support a multi-omic convergence on early mitochondrial and metabolic dysfunction localized to CA1.

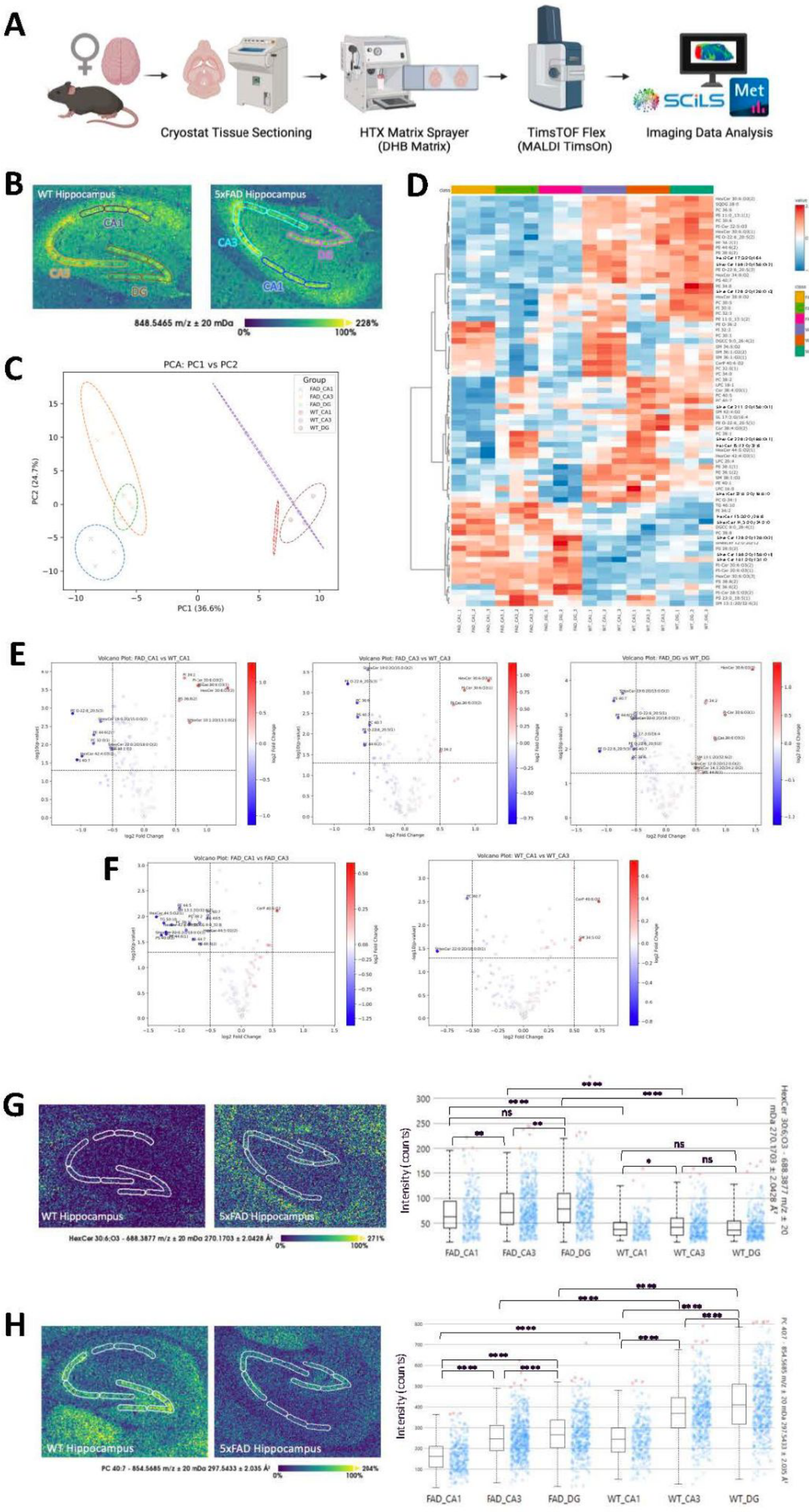
A
Cryostat Tissue Sectioning
HTX Matrix Sprayer
(DHB Matrix)
TimsTOF Flex
(MALDI TimsOn)
SCiLS
Met
Imaging Data Analysis
B
WT Hippocampus
5xFAD Hippocampus
CA1
CA3
DG
848.5465 m/z ± 20 mDa
0%
100%
228%
C
PCA: PC1 vs PC2
Group
FAD_CA1
FAD_CA3
FAD_DG
WT_CA1
WT_CA3
WT_DG
PC1 (36.6%)
PC2 (24.7%)
D
E
Volcano Plot: FAD_CA1 vs WT_CA1
Volcano Plot: FAD_CA3 vs WT_CA3
Volcano Plot: FAD_DG vs WT_DG
log2 Fold Change
F
Volcano Plot: FAD_CA1 vs FAD_CA3
Volcano Plot: WT_CA1 vs WT_CA3
G
WT Hippocampus
5xFAD Hippocampus
HexCer 30:6;O3 - 688.3877 m/z ± 20 mDa 270.1703 ± 2.0428 Å²
0%
100%
271%
Intensity (counts)
ns
FAD_CA1
FAD_CA3
FAD_DG
WT_CA1
WT_CA3
WT_DG
H
WT Hippocampus
5xFAD Hippocampus
PC 40:7 - 854.5685 m/z ± 20 mDa 297.5433 ± 2.035 Å²
0%
100%
284%
Intensity (counts)
FAD_CA1
FAD_CA3
FAD_DG
WT_CA1
WT_CA3
WT_DG

***Supplemental Figure 1.*** *Mass spectrometry imaging data illustrates lipid differences within hippocampal regions in 1 month old 5xFAD mouse. (A) MALDI-MSI (matrix assisted laser desorption/ionization – mass spectrometry imaging) workflow for brain tissue analysis (created with the BioRender scientific illustration software). Female snap frozen brain tissue of 1 month old 5xFAD mouse and WT counterpart were sectioned horizontally into 12 µm-thick slices at -23 °C using a cryostat (Leica CM3050 S). Horizontal sections exposing the hippocampus region of interest were adhered to ITO slides (Bruker MALDI IntelliSlides), with comparative 5xFAD and WT sections on one slide. One slide (N=1) was collected, dried in a vacuum desiccator for 30 min, then stored at -80 °C until use. Before analysis, tissues were thawed in a vacuum desiccator for 15 min at room temperature. DHB matrix (40 mg/mL) was prepared in 70% MeOH in water and deposited onto the slide using the HTX M3+ Sprayer. Spraying parameters were 75 °C temperature, 10 psi pressure, 100 µL/min flowrate, 1200 mm/min velocity, 10 second drying time, and 8 passes. MSI data was collected on the TimsTOF flex mass spectrometer (Bruker Scientific, LLC, Bremen, Germany). The laser was set to 20 µm diameter and collected over a mass range of 100-1300 Da under positive ionization mode. The raw imaging data was processed in Bruker's SCiLS Lab software for analysis and root mean square (RMS) normalization. Lipid MSI peaks were selected using 75% T-ReX[3] feature finding and exported for annotation in Metaboscape with lipid species and MS-DIAL spectral library[1] based on exact mass and collision cross section (CCS). (B) MSI data in SCiLS Lab resolved hippocampal subregions at m/z 849.5527. Based on this ion image, the CA1, CA3, and DG regions of WT and 5xFAD were segmented into three smaller subregions. The RMS normalized, average peak area intensity for all 18 subregions were generated and exported out of SCiLS for data visualization with Python. Prior to Python, the data was normalized using MetaboAnalyst 6.0 (low-abundance filtering by 10% of mean intensity value, normalization by sum, and log2 data transform). (C) Principal component analysis (PCA) plot showed separation of 5xFAD and WT hippocampus lipid* *profiles along PC1 and separation of hippocampal subregions along PC2. Hippocampal subregion separated along PC2 similarly between genotypes. (D) Heatmap of lipids, generated in MetaboAnalyst after ANOVA, show top 75 significantly expressed lipids between genotypes with hierarchical clustering of lipid features (Euclidean distance similarity measure and Ward's linkage). (E) Volcano plots showing differentially expressed lipids between subregions (eg. FAD_CA1 vs WT_CA1). Double filtering was set to include |FC| > 0.5 and p-value < 0.05. Labeled dots in red represent lipids increased in Hets after double filtering, and labeled dots in blue represent lipids decreased in 5xFAD. (F) Volcano plots showing differentially expressed lipids between same genotype subregions. Labeled dots in red represent lipids increased in CA1 after double filtering, and labeled dots in blue represent lipids decreased in CA1. (G) From the volcano plots, a shared upregulated lipid in 5xFAD hippocampal subregions was identified as HexCer 30:6;O3 [M+K]+. The ion image validates external data analysis results, visualizing a higher intensity signal in the 5xFAD subregions. The corresponding ion intensity box plots generated in SCiLS Lab confirmed higher abundance of HexCer 30:6;O3 [M+K]+ in 5xFAD subregions. (H) Across both genotypes, a shared downregulated lipid in the CA1 subregion compared to the CA3 subregion was identified as PC 40:7 [M+Na]+. The corresponding ion intensity box plots generated in SCiLS Lab further confirmed lower abundance of PC 40:7 [M+Na]+ in the CA1 subregion across genotypes, but also an overall lower abundance in 5xFAD hippocampus. P-values were assessed in SciLS Lab using the two-sided t test: ns ≥ 0.05, * P < 0.05, ** P < 0.01, *** P < 0.001, **** P< 0.0001. (Tsugawa et al., 2015).*

frontiers
